# The allotetraploid origin and asymmetrical genome evolution of common carp *Cyprinus carpio*

**DOI:** 10.1101/498774

**Authors:** Peng Xu, Jian Xu, Guangjian Liu, Lin Chen, Zhixiong Zhou, Wenzhu Peng, Yanliang Jiang, Zixia Zhao, Zhiying Jia, Yonghua Sun, Yidi Wu, Baohua Chen, Fei Pu, Jianxin Feng, Jing Luo, Hanyuan Zhang, Hui Wang, Chuanju Dong, Wenkai Jiang, Xiaowen Sun

**Affiliations:** Centre for Applied Aquatic Genomics, Chinese Academy of Fishery Sciences, Beijing, 100141, China; State Key Laboratory of Marine Environmental Science, College of Ocean and Earth Sciences, Xiamen University, Xiamen 361102, China; Novogene Bioinformatics Institute, Beijing, 100029, China; laboratory for Marine Biology and Biotechnology, Pilot National Laboratory for Marine Science and Technology, Qingdao, 266071, China; State Key Laboratory of Large Yellow Croaker Breeding, Ningde Fufa Fisheries Company Limited, Ningde, 352130, China; Heilongjiang River Fisheries Research Institute, Chinese Academy of Fishery Sciences, Harbin, 150001, China; Key Laboratory of Biodiversity and Conservation of Aquatic Organisms, Institute of Hydrobiology, Chinese Academy of Sciences, Wuhan, 430072, China; Henan Academy of Fishery Sciences, Zhengzhou, 450044, China; State Key Laboratory for Conservation and Utilization of Bio-Resources in Yunnan, Yunnan University, Kunming, 650091, China; College of Fisheries, Henan Normal University, Xinxiang, Henan 453007, China

## Abstract

Common carp (*Cyprinus carpio*) is an allotetraploid Cyprinid species derived from recent whole genome duplication and provides an excellent model system for studying polyploid genome evolution in vertebrates. To explore the origins and consequences of tetraploidy in *C. carpio*, we generated three chromosome-level new reference genomes of *C. carpio* and compared them to the related diploid Cyprinid genome sequences. We identified a progenitor-like diploid Barbinae lineage by analysing the phylogenetic relationship of the homoeologous genes of *C. carpio* and their orthologues in closely related diploid Cyprinids. We then characterized the allotetraploid origin of *C. carpio* and divided its genome into two homoeologous subgenomes that are marked by a distinct genome similarity to their diploid progenitor. On the basis of the divergence rates of homoeologous genes and transposable elements in two subgenomes, we estimated that the two diploid progenitor species diverged approximately 23 million years ago (Mya) and merged to form the allotetraploid *C. carpio* approximately 12.4 Mya, which likely correlated with environmental upheavals caused by the extensive uplift of the Qinghai-Tibetan Plateau. No large-scale gene losses or rediploidization were observed in the two subgenomes. Instead, we found extensive homoeologous gene expression bias across twelve surveyed tissues, which indicates that subgenome B is dominant in homoeologous expression. DNA methylation analysis suggested that CG methylation in promoter regions plays an important role in altering the expression of these homoeologous genes in allotetraploid *C. carpio*. This study provides an essential genome resource and insights for extending further investigation on the evolutionary consequences of vertebrate polyploidy.

## INTRODUCTION

Two rounds of whole genome duplication (2R WGD) occurred during the evolution of early vertebrates before the divergence of lamprey from jawed vertebrates. An additional round (3R) of whole genome duplication occurred in ray-finned fishes at the base of the teleosts^1–4^. The 3R WGD, which is also known as the teleost-specific WGD (Ts3R), was estimated to happen approximately 320 million years ago (Mya)^5^. The duplication of entire genomes plays a significant role in evolution. Multiple rounds of WGD produced redundant genes, which provided an important genetic material basis for phenotypic complexity, which would potentially benefit an organism in its adaptation to environmental changes^6^. Beyond these WGD events, some teleost lineages encountered recent additional genome duplications and polyploidization. Most of the well-characterized and well-recognized polyploidy fishes are in Salmonidae^7,8^ and Cyprinidae^9–12^ (**Figure 1**). There was apparently only one autotetraploidization event that occurred at the common ancestor of salmonids approximate 80 Mya, while polypoloidization evolved independently on multiple occasions in Cyprinids, of which the common carp (*Cyprinus carpio*) and goldfish (*Carassius* sp.) appear to have experienced the latest allotetraploidization event before their divergence^13–15^, thus providing an excellent model system for investigating the initial allopolyploidization event in teleosts and understanding the evolutionary benefits for phenotypic plasticity, environmental adaptations and species radiation post the latest WGD. As one of the most important food and ornamental fishes in the Cyprinidae family, *C. carpio* has been widely cultured worldwide, with an annual production of over 4 million metric tons^16,17^. Owing to its importance in aquaculture and genome evolution studies, many efforts were made to develop genetic and genome resources in the past decades. Although the next-generation sequencing technologies and assembly algorithms have overcome many major obstacles for whole genome sequencing and assembly, the allotetraploid nature and highly heterozygous level of *C. carpio* genome still bring many challenges. Unlike diploid genomes, polyploid genomes involve multiple rounds of WGD and segmental duplications that harbour much more complex structures and gene contents than those of diploid genomes, which poses significant challenges in discriminating among homoeologous sequences and in producing high-quality genome assemblies. The first allotetraploid carp genome of its European subspecies *C. carpio carpio* Songpu strain had been previously finished but only anchored 52% of the scaffolds onto the 50 chromosomes^13^. The scaffolds and genes on approximately half of the genome remained ambiguous with respect to homoeologous relationships, thus creating a great obstacle for investigating the allotetraploid genome evolution of carps.

**Figure 1.**
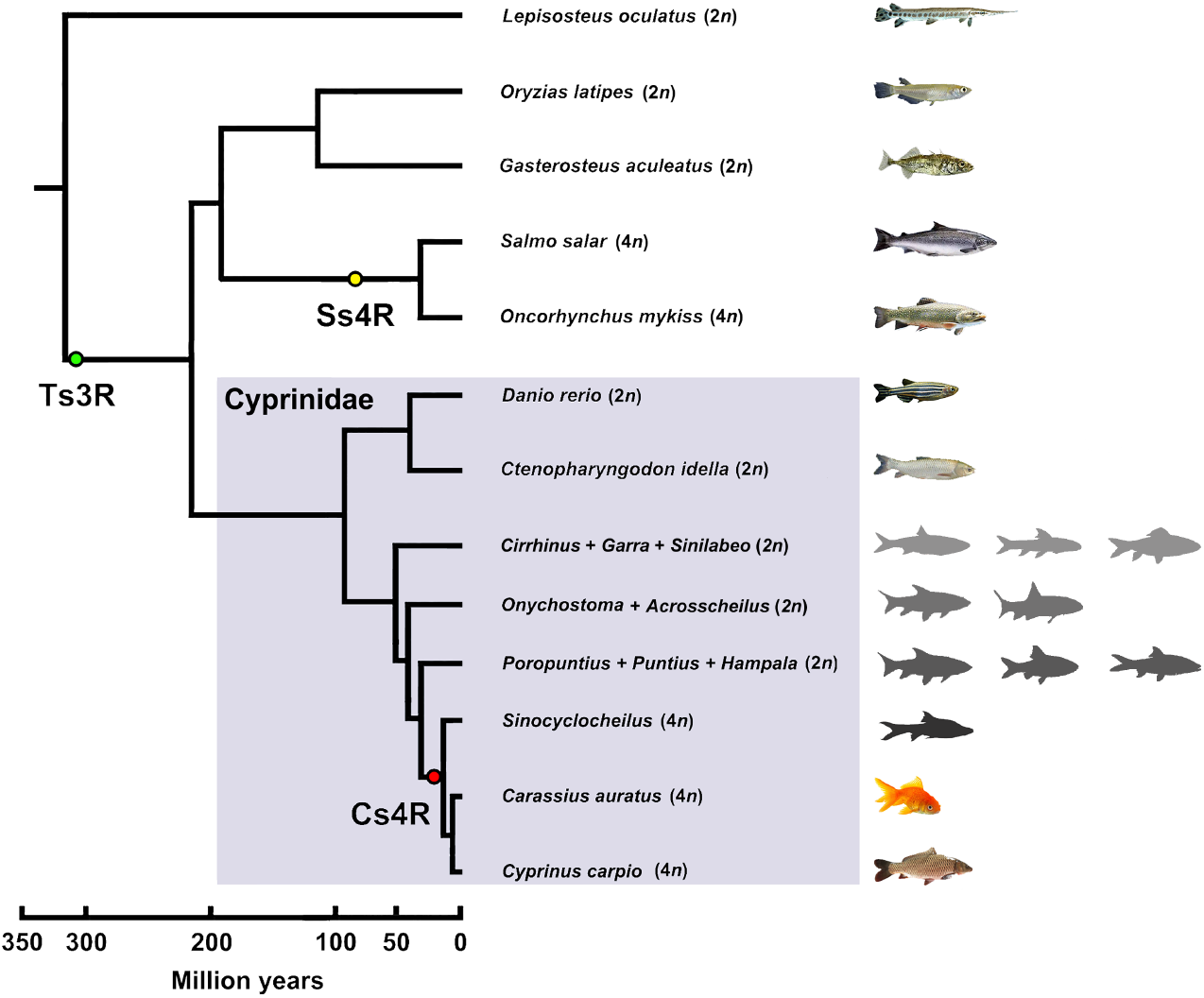
Phylogenetic relationship of tetraploid Cypinidae and relevant teleost lineages. The phylogenetic topologies and divergence ages are taken from the TimeTree database (ref. 11) and ref. 12. Green, yellow and red circles represent the teleost-specific whole genome duplication (Ts3R), salmonid-specific whole genome duplication (Ss4R) and carp-specific whole genome duplication (Cs4R), respectively.

To better understand the tetraploid genome structure and gain insights into the post-WGD evolution of *C. carpio*, it is vital to obtain allotetraploid genomes with higher accuracy and connectivity. We therefore sequenced and assembled chromosome-level allotetraploid genomes of three different *C. carpio* subspecies. Moreover, we also sequenced four closely related diploid Cyprinid genomes and identified potential ancestral diploid lineages to discriminate between two highly similar subgenomes.

The post-WGD evolution of the newly merged allotetraploid vertebrate genome has been investigated for its evolutionary history, subgenome structure and content, differentiated selective pressure, asymmetrically expressed homoeologous genes and their epigenetic regulations. Together, these resources and findings on the allotetraploid genome evolution of *C. carpio* provide a foundation for further unveiling the genetic basis of polyploid complexity, adaptation and phenotypic advantages and for accelerating the genetic improvement of those polyploid fishes for aquaculture.

## RESULTS

### Genome assembly, annotation and scaffold anchoring

We sequenced the tetraploid genomes of three distinct *C. carpio* strains, namely, Hebao red carp (HB) and Yellow River carp (YR) from China, which belong to the subspecies *C. carpio haematopterus*, and German mirror carp (GM) from Europe, which belongs to the subspecies *C. carpio*, by whole-genome shotgun methods (**Supplementary Figure 1, Supplementary Table 1**). The three assembled genomes spanned 1,460 Mb, 1,425 Mb, and 1,416 Mb, with the contig N50 and scaffold N50 of 20.68 kb and 923.37 kb for HB, 21.81 kb and 1,706 kb for YR, and 52.14 kb and 3,466 kb for GM, respectively (**Supplementary Table 2**). We constructed three high-resolution genetic maps with 29,019 (HB), 28,194 (YR) and 32,160 (GM) markers on 50 chromosomes by genotyping the mapping families using a carp 250K SNP array^18^ (**Supplementary Figure 2, Supplementary Tables 3**). We then anchored the assembled genomes to the 50 chromosome frames of the genetic maps. Finally, three chromosome-level reference genomes of *C. carpio* were created with high connectivity, representing 1.24 Gb (82%) of HB, 1.26 Gb (89%) of YR and 1.3 Gb (92%) of GM assemblies, respectively (**Supplementary Table 4**). All three assemblies represent a substantial improvement over the previously published draft genome sequence of *C. carpio*, which only anchored 875 Mb (52%) onto the 50 chromosomes^13^.

We BLAST-aligned highly conserved core eukaryotic genes (Cluster of Essential Genes (CEG) database) to the genome assemblies with a core eukaryotic genes mapping approach (CEGMA) pipeline, which showed high-confidence hits identified in three assembled genomes of *C. carpio* (**Supplementary Table 5**). We also validated the assembled genomes by matching them with expressed sequence tags (ESTs) downloaded from the US National Center for Biotechnology Information (NCBI) database, which indicated that 99.86%, 99.26% and 99.20% of the ESTs were covered by the assembled genomes of HB, YR and GM, respectively (**Supplementary Table 6**). We further compared the assembled genome of GM with a previously published draft genome of the mirror carp Songpu strain (SP). Both the GM and SP belong to the European subspecies *C. carpio carpio*. To assess the genome connectivity and assembly accuracy, we aligned 34,932 mate-paired BAC-end sequences (BES) that were derived from the SP genome to both SP and GM assemblies. The result showed that 98% and 97% of the BESs were mapped to the genomes of SP and GM, respectively. In over 85% of BES mate-pairs, both BESs were aligned to the same scaffold of GM, compared to only 31% of the BES mate-pairs that aligned to the scaffold of the SP draft genome. We observed the standard Poisson distribution of the BES pair intervals on GM scaffolds that corresponded to the real BAC insertion length, while no regular distribution was observed on the SP scaffolds (**Supplementary Figure 3**). A comparative analysis of three *C. carpio* genome assemblies revealed that the three genomes are highly conserved at the chromosome level, with only a limited number of large structures or segmental variations (**Supplementary Figure 4**). We identified 488.15 Mb of repetitive sequences from the *HB* genome, 449.18 Mb from the YR genome and 428.29 Mb from the GM genome, which contributed to 31.52%, 32.32% and 30.24% of three genomes, respectively, with similar classifications and proportions (**Supplementary Table 7**). The most abundant transposable elements were DNA transposons, which contributed to approximately 13% of all three genomes, with Tc1 mariners representing approximately 5% of the genomes. We annotated 44,269, 44,626 and 44,758 protein-coding genes in the HB, YC and GM genomes, respectively (**Supplementary Table 8**). Approximately 96.9%, 96.2% and 96.1% of HB, YC and GM genes could be annotated by non-redundant nucleotides and proteins in the SWISS-PROT, Gene Ontology (GO), Kyoto Encyclopedia of Genes and Genomes (KEGG), Cluster of Orthologous Groups (COG), Pfam and NCBI databases (**Supplementary Table 9**). Together, the evidence suggested that the three newly assembled genomes of *C. carpio* had been improved significantly in connectivity and contiguity, thus paving the way for further unveiling of the allotetraploid genome evolution of Cyprinids.

### Allotetraploid origin of the common carp *C. carpio*

The common carp *C. carpio* resulted from the ancient hybridization of two ancestral diploid cyprinid species^1^, which is of critical importance for genome evolution studies that divide the allotetraploid genome into two subgenomes, thereby representing two ancestral diploid genomes. Cyprinids are a diverse teleost family with over 2,400 valid species in at least 220 genera^19,20^, of which the subfamily Cyprininae comprises over 1,300 species in four groups of barbine, cyprinine, labeonine and schizothoracine with diversified and complex karyotypes from 2n = 50 to ~470^12^. Under such circumstances, it is very challenging to identify diploid ancestral lineages and unveil the evolutionary origin of allotetraploid common carp. Previously, we successfully recognized 25 homoeologous chromosome pairs in the *C. carpio* genome by aligning 50 chromosomes of *C. carpio* into 25 chromosomes of the zebrafish (*Danio rerio*) diploid genome^13,21,22^, thereby revealing the existence of two homoeologous sets of chromosomes in the *C. carpio* genome. To explore the evolutionary relationship of *C. carpio* and its closely related tetraploid and diploid Cyprininae species, we constructed a phylogenetic tree of the representative Cyprininae species using a nuclear gene, *recombination activating gene 2* (*rag2*), which presents only one copy in diploid cyprinids but two copies in tetraploid cyprinids (**Figure 2a, Supplementary Figure 5, Supplementary Table 10**)^12,23^. The phylogenetic topology revealed that two copies of the *rag2* of the closely related tetraploid genomes, including three *C. carpio*, three cavefish *Sinocyclocheilus* and goldfish *Carassius auratus* genomes, were clustered into two distinct homoeologous clades, which suggested that these tetraploid cyprinids either derived from a common tetraploidization event and then diverged into different species or experienced independent and recurrent tetraploidization events involving the same or closely related diploid ancestors. One of the homoeologous clades of *rag2b* from the tetraploid genomes subsequently joined with *rag2* genes from a diploid group in Barbinae, including multiple genera, such as *Poropuntius*, *Puntius, Hampala* and *Onychostoma*, which implied that one of the two diploid progenitors of *C. carpio* may have derived from a diploid Barbinae species. However, the *rag2a* clade did not merge with any *rag2* genes from the diploid species. The phylogenetic results gave us clues on how to separate the two subgenomes of the allotetraploid *C. carpio* genome. We reasoned that the genome similarity and alignment coverage between the diploid progenitors and descendent allotetraploid on each homoeologous chromosome pair would facilitate the discrimination of two descendent subgenomes. We thus selected three species from Barbinae *(Poropuntius huangchuchieni, Hampala macrolepidota* and *Onychostoma barbatulum*) as progenitor-like diploid candidates and one species from a relatively distant lineage *(Cirrhinus molitorella*) as a reference diploid species for whole genome sequencing and draft assembly (**Supplementary Figure 6, Supplementary Table 11**). The genome sequences of the four newly sequenced diploid Cyprinids and two previously sequenced diploid Cyprinids (*Ctenopharyngodon idella* and *D. rerio*) were then aligned to 25 pairs of homoeologous chromosomes of *C. carpio* genomes. The results showed that the genome sequences of three diploid Barbinae species have an extraordinarily higher similarity and coverage in one chromosome than in the other for each homoeologous chromosome pairs without exception, while no significant similarity or coverage differences were observed when the genome sequences of *C. molitorella, C. idella* or *D. rerio* were aligned to the homoeologous chromosome pairs of *C. carpio* (**Figure 2b and 2c**). The results provided substantial evidence of the allotetraploid origin of *C. carpio* and suggested that one subgenome progenitor possibly originated from a diploid lineage of Barbinae, while the other progenitor might be an unexplored or even extinct diploid from a relatively distant lineage in Cyprinids. We therefore divided 50 chromosomes evenly into subgenomes A and B, which represented the unknown progenitor A and the ancient progenitor B from Barbinae, respectively. We further performed a phylogenomic analysis based on 2,071 conserved homoeologous gene pairs among two tetraploids (*C. carpio* and *S. anshuiensis)* and their orthologues from three diploids (*D. rerio, C. idella* and *P. huangchuchieni*) and confirmed the homoeologous relationship of two subgenomes in *C. carpio* (**Supplementary Figure 7**), thus underpinning that subgenome B of *C. carpio* indeed originated from a diploid Barbinae species in Southwest China. These findings built the foundation for studying the polyploid origination and evolution in *C. carpio* and its closely related species and appear promising for generating synthetic polyploid lines with their diploid progenitors to achieve the desired breeding purposes.

**Figure 2.**
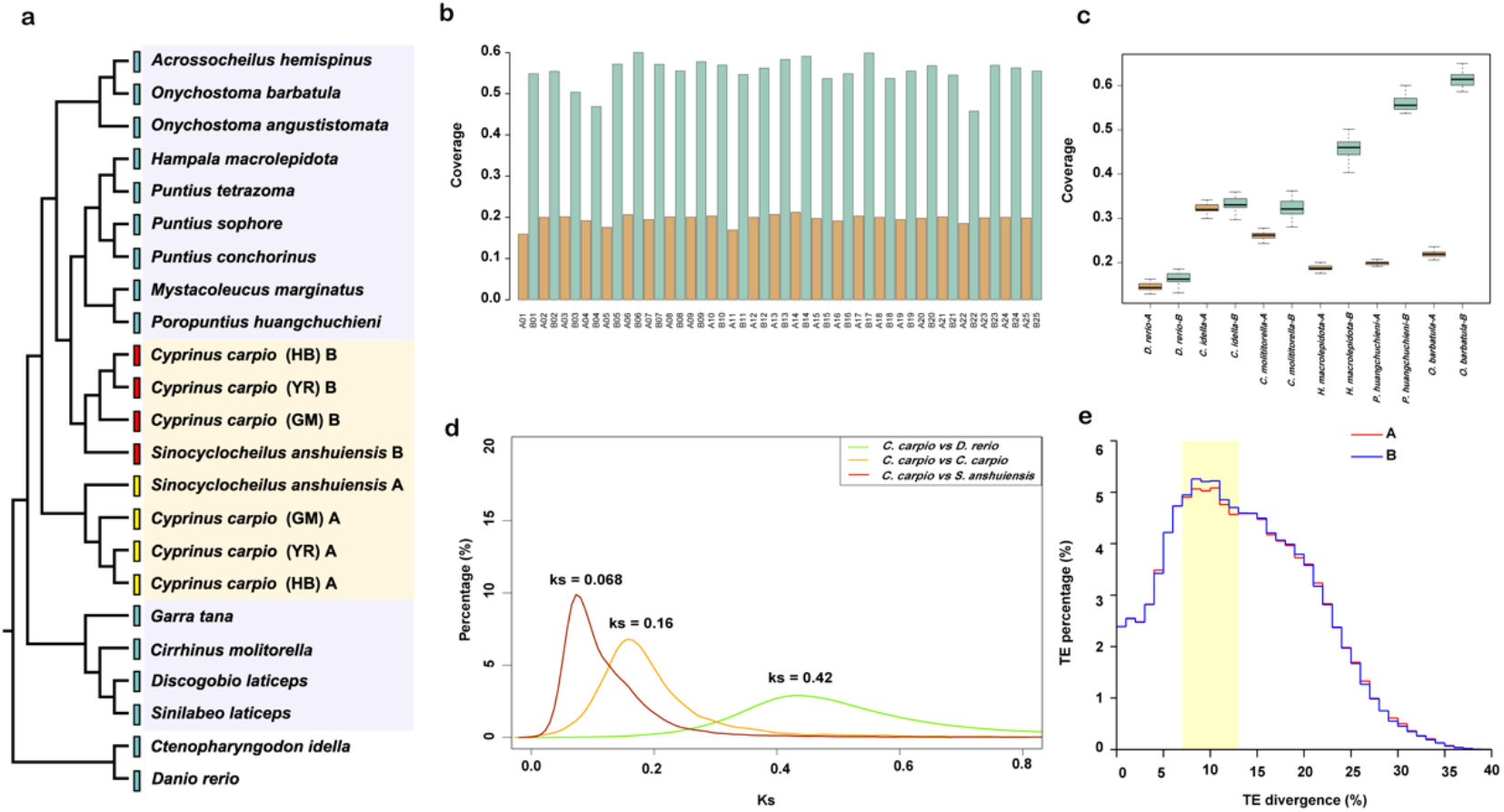
Allotetraploid origin and evolution history of *C. carpio*. **a.** Phylogenetic relationship of RAG2 orthologues of *C. carpio* and its tetraploid and diploid close relatives in the subfamily Cyprininae. The pentagrams indicate three selected diploid species (*Poropuntius huangchuchieni, Hampala macrolepidota*, and *Onychostoma barbatulum)* as progenitor-like diploid candidates, and one species from a relatively distant lineage (*Cirrhinus molitorella)* for genome sequencing to represent the closely related diploid lineages from Cyprininae. **b.** A histogram shows the coverage of *P. huangchuchieni* genome sequence mapping to 50 chromosomes of *C. carpio*. Chromosome IDs have been re-assigned to represent two sets of homoeologous chromosomes, **c.** Boxplots show the genome coverage and similarity comparisons of diploid relatives to the tetraploid genome of *C. carpio*. **d.** The distribution of the synonymous substitution rates (Ks) of homologous genes between I), *danio* and *C. carpio*, *C. carpio* and *Sinocyclocheilus*, and homoeologous genes between two subgenomes of *C. carpio*. Three peaks (Ks = 0.42, 0.16 and 0.068) of Ks distribution indicate the divergences of *D. danio* and *C. carpio*, *C. carpio* and *Sinocyclocheilus*, and two progenitors of *C. carpio*. **e.** The distribution of sequence divergence rates of transposable elements (TEs) as percentages of subgenome sizes of *C. carpio*. The TE content segregation between subgenomes A and B indicates the events of diploid progenitor divergence and subgenome merger.

To estimate the accurate time of the Cyprinid-specific allotetraploidization event, we calculated the synonymous substitution rates (Ks) of 8,270 homoeologous genes to determine the divergence time of the two subgenomes. These homoeologous genes present only one copy in the diploid genome of *D. rerio* and one copy in each of the two subgenomes in the allotetraploid genome of *C. carpio*. The substitution rates of *Danio-Cyprinus* paralogous genes, the homoeologous genes in subgenomes A and B in *C. carpio*, and the *Cyprinus-Sinocyclocheilus* paralogous genes were calculated to be 0.42, 0.16 and 0.068, respectively (**Figure 2d**). We applied the previously determined molecular clock that Ks in teleost is approximately 3.51 × 10^−9^ substitutions per synonymous site per year and therefore estimated that *D. rerio* and *C. carpio* diverged approximately 60 million years ago (Mya), the two ancient progenitor species of *C. carpio* diverged approximately 23 Mya, and *Sinocyclocheilus* and *Cyprinus* diverged approximately 9.7 Mya. Thus, we estimated that the Cyprinid-specific WGD and allotetraploidization event most likely occurred after the divergence of two ancient progenitors (23 Mya) but before the divergence of the two allotetraploid lineages of *Sinocyclocheilus* and *Cyprinus* (9.7 Mya). We further collected transposable elements (TE) from two subgenomes and assessed their divergence rates in each subgenome. Intriguingly, we identified differentiated TE contents in subgenomes A and B with divergence rates from 7% to 13%, which formed a “bubble” peak in the TE divergence profile (**Figure 2e**). This suggested that TE substitution rates in subgenomes A and B had to be differentiated after their divergence into two diploid progenitors (23 Mya) and were re-unified in the allotetraploid genome after the Cc4R event. We therefore estimated that the WGD event likely occurred approximately 12.4 Mya based on the TE substitution rate, which was consistent with a previous estimation based on microsatellite loci^1^ but much earlier than the estimation based on the previous draft genome^13^. The estimated WGD time was in the middle Miocene and correlated with the extensive uplift of the Qinghai-Tibetan Plateau (QTP) (**Supplementary Figure 8**). The air circulation was significantly altered by the QTP uplift event, together with the global cooling in the middle Miocene, and contributed to massive cooling and aridification in Central Asia^24^. Environmental upheavals have been proposed as one of the major driving forces of the WGD and polyploidization^25,26^. We reasoned that the latest occurring WGD conferred distinct adaptive advantages to polyploidized *C. carpio*, which allows them to survive and thrive in a drastically changed environment that poses challenges to those diploid progenitors and relatives.

### Subgenome structure and gene content

We divided the allotetraploid *C. carpio* genome evenly into two distinct subgenomes, A and B, with 25 chromosomes in each based on the differentiated sequence similarities in comparison with the genome of progenitor B from Barbinae. The new chromosome IDs were then assigned to 50 chromosomes based on their homologous relationship with 25 chromosomes of *D. rerio* (**Supplementary Table 12**). For instance, chromosomes 1A and 1B were assigned to a pair of homoeologous chromosomes that are syntenically related to chromosome 1 of *D. rerio*, of which 1A belongs to subgenome A and 1B belongs to subgenome B. The new chromosome IDs will facilitate better understanding of the homoeologous landscape of the allotetraploid *C. carpio* genome.

Previous studies on allopolyploid genomes, mostly in plants, revealed that one of the parental subgenomes often retains significantly more genes and exhibits significantly higher expression, stronger purifying selection and a lower DNA methylation level than those of the other subgenome. This phenomenon was referred to as subgenome dominance^27,28^. It is essential to investigate the homoeologous gene contents, the expression profile and their epigenetic regulations for verifying subgenome dominance and to better understand the allotetraploid genome evolution after genome merging. Therefore, we assigned 21,078 genes in 633 Mb sequences to the subgenome A and 22,099 genes in 671 Mb sequences to the subgenome B of *C. carpio*, indicating that subgenome B retains 1,021 (~5%) more genes and is slightly more dominant than subgenome A (**Supplementary Table 13**). The GC content, gene structure, and repetitive element distribution did not show significant differences between the two subgenomes (**Supplementary Figure 9, Supplementary Table 14**). To assess the fate of the homoeologous genes in the *C. carpio* genome, we compared the gene contents of the allotetraploid genome of *C. carpio* and the closely related diploid genome of *C. idella* and built a total of 10,724 orthologous gene pairs or triplets within the two genomes. We identified 8,291 orthologous gene triplets that presented one copy in the diploid *C. idella* genome and one copy in the homoeologous chromosome of each subgenome of the *C. carpio* genome. Visualization of the chromosomal locations of 8,291 homoeologous gene pairs revealed a high degree of chromosome-level synteny between the subgenomes A and B (**Figure 3a, Supplementary Figure 4**). We also found 915 and 1,220 single-copy orthologous genes in the subgenomes A and B of *C. carpio*, respectively, suggesting that unequal loss of the homoeologous genes from ancestral progenitor genomes had occurred in the two subgenomes (**Supplementary Table 15**). Extensive gene losses and rediploidization are commonly observed following polyploidy events; however, our result indicated that only a small portion (~4.9%) of duplicated genes had been lost after the latest WGD of *C. carpio* (**Supplementary Figure 12**). A gene ontology (GO) analysis indicated that gene loss was not random in the *C. carpio* genome after the latest WGD. Single-copy genes are overrepresented in some essential functional categories in both subgenomes, such as the nucleic acid metabolic process (GO:0090304), DNA repair (GO:0006281), DNA replication (GO:0006260), nuclease activity (GO:0004518) and ribonucleoprotein complex (GO: 1990904), which was consistent with previous studies in various tetraploid genomes^29–32^ (**Supplementary Table 16-23**) and suggested that these genes might be sensitive to altered gene dosage during WGD and might lose reciprocally to return single-copy status in two subgenomes. We found that only a small portion of homoeologous gene pairs (118 homoeologs) were mapped within the same subgenome, suggesting that limited homoeologous exchange events have occurred across two subgenomes after the WGD event (**Supplementary Table 24 and 25**). We identified a total of 92 large segmental rearrangements, including 40 segmental inversions, between the homoeologous chromosomes of two subgenomes. For example, we found a segmental inversion of 2.5 Mb and a segmental translocation of 1.8 Mb presenting in the homoeologous chromosomes A24/B24 and A15/B15 of the *C. carpio* genome, respectively (**Supplementary Figure 13**). The chromosome-level TE distribution analysis revealed that TEs are significantly enriched in the flanking regions of the structure variations between the two subgenomes, suggesting that TEs could be the potential driving force of those homoeologous exchanges and segmental variations in the allotetraploid genome of *C. carpio*. Thus, the gene content and syntenic analyses revealed that two homoeologous gene sets were well preserved despite minor gene losses in the two subgenomes of the allotetraploid *C. carpio* genome, and subgenome B retains slightly more genes than subgenome A.

**Figure 3.**
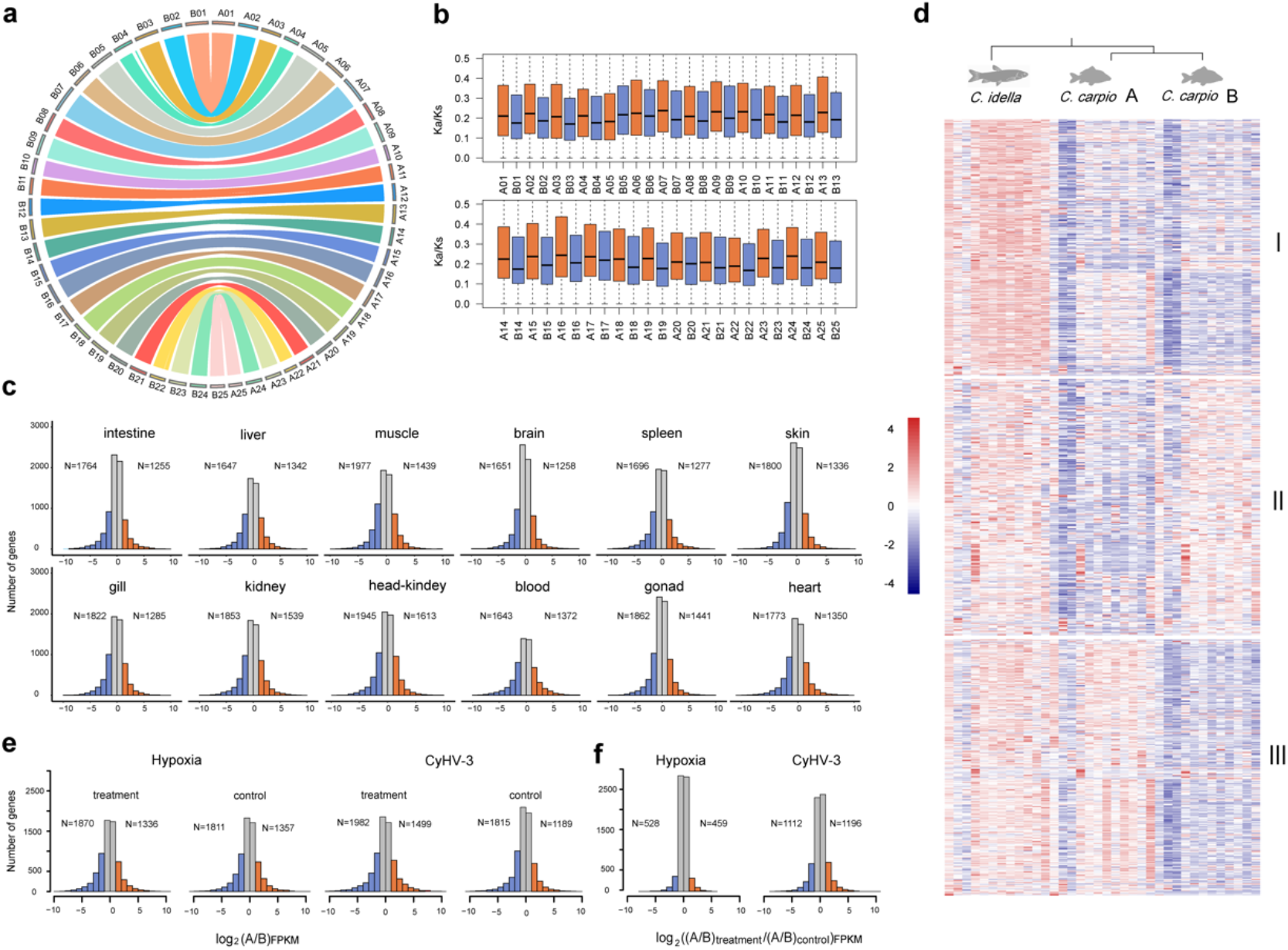
Asymmetrical homoeologous gene expression in *C. carpio*. **a.** Circos plot distribution of homoeologous gene pairs in 25 chromosome pairs across subgenomes **A** and B. **b.** Boxplot of the Ka/Ks ratio distribution of protein-coding genes in 50 chromosomes of *C. carpio*. Red and blue boxplots indicate that the chromosomes belong to subgenomes **A** and B, respectively, **c.** Histograms of the genome-wide expression of homoeologous genes among the indicated tissues of *C. carpio*. **N** values indicate the number of dominant genes in subgenomes **A** and B. d. Heatmaps of three divergently expressed triplet clusters (each triplet includes two homoeologous genes of *C. carpio* and their orthologue of *C. idella*), indicate potential subfunctionalization and neofunctionalization in the allotetraploid genome of *C. carpio*. e. Histograms of the genome-wide expression divergence of homoeologous genes of *C. carpio* in stress treatments and controls. **N** values indicate the number of dominant genes in subgenomes **A** and B, respectively, **f.** Histogram of the ratio of homoeologous expression divergence in the stress treatment and controls [(A/B)_treatment_/(A/B)_control_], which indicated accelerated homoeologous expression divergence in stress treatments compared to controls.

### Subgenome expression bias in the allopolyploid *C. carpio* genome

Subgenome dominance usually leads to relaxed purifying selection and the dominant gene expression of homoeologous genes in many tetraploid genomes of plants and animals^28,33–35^. To assess the selective pressure of two subgenomes, we calculated both the nonsynonymous substitution rate (Ka) and Ks values based on homoeologous gene pairs (**Supplementary Table 26**). Intriguingly, the results showed that all 25 chromosomes in subgenome A had a significantly higher Ka/Ks ratio (mean Ka/Ks = 0.20) than their homoeologous chromosomes in subgenome B (mean Ka/Ks = 0.18) (*p* = 1.28e-15). The distinct selective pressure difference of 25 paired homoeologous chromosomes indicated asymmetric evolution of two subgenomes in *C. carpio*. The subgenome A genes are under stronger positive selection and are evolving faster than their homoeologs in subgenome B. The subgenome B genes are under greater purifying selection than their homoeologs in subgenome A (**Figure 3b**). A previous study on the tetraploid maize genome suggested that dominant gene expression is a strong determinant of the strength of purifying selection^36^. To investigate whether the similar phenomena also presents in the *C. carpio* genome, we therefore compared the genome-wide transcriptional levels of subgenomes A and B based on the gene expression levels of 8,291 homoeologous gene pairs in 12 tissues to investigate the homoeologous gene expression patterns and their divergence in two subgenomes (**Figure 3c, Supplementary Table 27**). The results indicated that a total of 7,536 expressed homoeologous gene pairs (~91%) had expression differences greater than 2-fold change, including 4,719 and 5,403 homoeologs having higher expression in subgenomes **A** and **B** in at least one tissue, respectively (**Supplementary Table 28-51**), of which, 2,133 and 2,817 homoeologs had higher expression values exclusively in the 12 tissues of subgenomes A and B, respectively, while 2,586 homoeologs had swinging expression bias in 12 tissues. The homoeologous expression bias showed asymmetric expression patterns between subgenomes A and B so that the genome-wide expression level dominance was biased towards subgenome B in the allotetraploid *C. carpio* genome (**Supplementary Figure 14, Supplementary Table 52**). We tuned the threshold of expression change to identify those homoeologs with extraordinary expression divergence in the two subgenomes. We found that 1,018 homoeologous genes had expression differences greater than 32-fold between the two subgenomes, including 406 and 627 homoeologs that had higher expression exclusively in subgenomes A and B, respectively. Gene annotation indicated that the dominantly expressed homoeologs of subgenome A were enriched in “nucleobase biosynthetic and metabolic processes”, “lipid metabolic process” and “lipid biosynthetic process”, while dominantly expressed homoeologs of subgenome B were enriched in “oxidoreductase activity”, “hydrolase activity”, “response to stress” and “DNA repair” (**Supplementary Table 53-58**).

We further built co-expression clusters across all 12 tissues to assess the overall expression divergence rate between the two subgenomes (**Supplementary Figure 15**). Of the 8,291 homoeologs in eight co-expression clusters, 1,986 pairs of homoeologous genes (~24%) had been assigned to different co-expression clusters. The results suggested that substantial spatial partitioning of the homoeologous differentially expressed genes occurred, while the majority of the homoeologous genes still tend to retain similar expression levels and patterns in the allotetraploid genome of *C. carpio*. The spatially differentially expressed homoeologous gene pairs in two subgenomes may have experienced or been experiencing functional divergence via the subfunctionalization or neofunctionalization mechanism, which are commonly observed in allopolyploid genomes after genome mergers and gene duplications. To discriminate which functional divergence mechanism potentially occurred on specific homoeologs, we further collected transcriptomic data from the same 12 tissues of *C. idella* (**Supplementary Table 59**), which was the most closely related diploid Cyprinid with a complete reference genome available and could serve as an orthologous reference of the 8,291 homoeologous gene pairs in *C. carpio* for an expression divergence analysis. We built co-expression clusters based on 8,214 expressed orthologous triplet genes in *C. idella* and *C. carpio*. The expression patterns of 306 orthologous triplets were differentially expressed in *C. idella* and in two subgenomes of *C. carpio*, suggesting that the subfunctionalization mechanism potentially re-shaped the spatial expression patterns of these homoeologs of *C. carpio*. We further identified that 293 genes in subgenome A and 228 genes in subgenome B had conserved co-expression patterns with their orthologues of *C. idella*, respectively, while their homoeologous copies in the opposite subgenome were differentially expressed, suggesting that neofunctionalization mechanism potentially occurred in one of the copies (**Figure 3d**). In addition, we also found that 191 and 620 homoeologous gene pairs were solely transcribed in subgenomes A and B, respectively, while the other copies were barely transcribed in 12 tissues, indicating that nonfunctionalization may have occurred and silenced one copy of the homoeologous pairs (**Figure 4c, Supplementary Table 60 and 61**). Selection pressure on these 811 homoeologs with an extreme divergent expression in two subgenomes was then investigated, which showed that the dominantly transcribed homoeologous copies, regardless of their subgenome location, were more likely experiencing stronger purifying selection than their homoeologs (**Figure 4d**). The observed connection of strength of purifying selection and expression dominance was consistent with previous findings in polyploid plants^28^. We observed that a total of 1,638 homoeologs, accounting for approximately 20% of surveyed homoeologous genes, showed spatial expression pattern divergence, suggesting that a majority of homoeologous genes still maintain conserved expression patterns and gene functions in the tetraploid *C. carpio* genome. We further investigated the gene functions of these functionally divergent genes or silenced genes in either subgenome A or B and found that some vital functional categories of GO and KEGG were disproportionately over-represented, including “fatty acid metabolism (KO01212)”, “RNA degradation (KO03018)”, “ribosome biogenesis (KO03008)”, “nucleotide excision repair (KO03420)”, “peroxisome (KO04146)”, “lipid metabolic process (GO:0006629)”, “nucleic acid metabolic process (GO:0090304)”, and “oxidation-reduction process (GO:0055114)”, which are potentially involved in critical pathways and require optimal stoichiometry of gene expressions to ensure normal biological processes and the survival of tetraploid *C. carpio* after genome merging (**Supplementary Table 62-65**). For example, we found that the eukaryotic translation initiation factor 3 subunit I (eIF3I) gene located on chromosome A19 is dominantly expressed in all 12 surveyed tissues, while its homoeologous copy on chromosome B19 is completely silenced (**Figure 4f**). eIF3 is a multiprotein complex that functions during the initiation phase of eukaryotic translation^37^. Altered expression levels of eIF3 subunits correlate with neurodegenerative disorders and cancer development and may also trigger infection cascades^38^. We also found that the expression of programmed cell death 1 ligand 1 (Pd-11) gene on chromosome B13 is suppressed in all 12 tissues, while its homoeolog on chromosome A13 maintains normal expression (**Supplementary Figure 16**). The Pd-11 gene has near-ubiquitous expression in tissues and plays a major role in suppressing the immune system by dampening the T cell response and preventing overactivation during proinflammatory states^39,40^. Similarly, the long-chain fatty acyl-CoAligase 6 (Acsl6) gene in chromosome B19 of subgenome B is silenced when its homoeolog on chromosome A19 of subgenome A is dominantly expressed (**Supplementary Figure 16**). The Acsl6 gene is mainly expressed in neural cells and the brain and is essential for regulating the partitioning of acyl-CoA species towards different metabolic fates, such as lipid synthesis or β-oxidation^41^. These genes are likely involved in dosage-sensitive regulation pathways by which the suppressed expression of one homoeologous copy would ensure optimal RNA and protein supplies and maintain normal biological processes.

**Figure 4.**
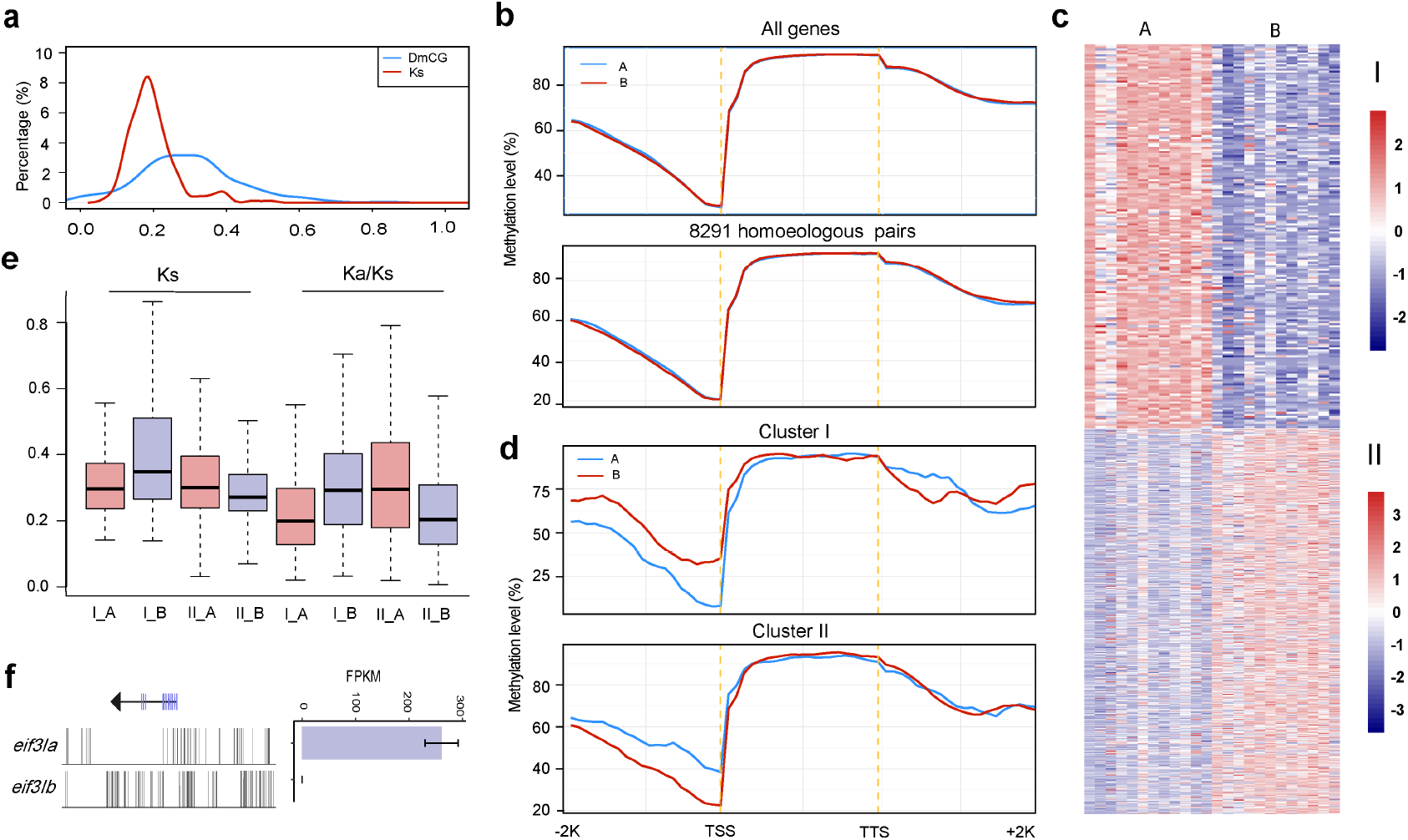
DNA methylation in homoeologous expression bias in *C. carpio*. **a.** Distribution of synonymous substitution values (Ks) (red) and gene-body DmCG percentages (blue) of 2,393 methylated homoeologous genes. Peak values are indicated by arrows, **b.** Heatmaps of two extremely divergent expression clusters, of which one of two homoeologous genes in subgenomes A or B was completely silenced in all 12 tissues, **c.** CG methylation levels of all annotated genes and 8,291 homoeologous gene pairs in two subgenomes, **d.** CG methylation levels of divergent expressed homoeologous genes corresponding to clusters I and II in **a. e.** Boxplot of the Ka/Ks ratio distribution of homoeologous genes of two extremely divergent expression clusters in two subgenomes as shown in **c.** Red and purple boxplots indicate the genes in subgenomes A or B, respectively, **f.** CG methylation level and expression level of homoeologous genes *eif3* of *C. carpio*, which demonstrated that reduced CG methylation levels in the promotor region correlated with increased expression levels of *eif3a*, while *eif3b* was heavily methylated and silenced. RPKM reads per kilobases per million.

Polyploidization may confer a significant adaptive advantage in response to various environmental challenges during the evolution history^26^. To explore whether homoeologous gene expression bias plays a role in stress responses, we collected transcriptome expression data from *C. carpio* under hypoxia and disease stresses with Cyprinid herpesvirus 3 (CyHV-3) and the bacterial pathogen *Aeromonas hydrophila* separately, as well as data from control samples from the same experiments. We investigated homoeologous expression divergence in 8,291 previously identified homoeologous gene pairs in both stress treatments and controls. We still observed asymmetric homoeologous expression patterns in all surveyed samples of biotic or abiotic stress treatments and their controls and found that subgenome B was still dominantly expressed (**Figure 3e, Supplementary Figure 17, Supplementary Table 66**). We also found that samples under stress treatment tend to have more homoeologs that are divergently expressed in two subgenomes than the control samples (**Figure 3f**) We performed differential gene expression (DEG) analysis of the homoeologous genes and identified 3,367 (23.47%), 5,768 (38.70%), and 3,448 (23.67%) genes that were differentially expressed with 2-fold changes compared with corresponding controls in the stress experiments with hypoxia, CyHV-3 and *A. hydrophila* stresses, respectively (**Supplementary Table 67**). To evaluate the homoeologous expression responses to specific stress treatments within the allotetraploid genome, we calculated the summed expressions of homoeologous genes in both treatments and controls, which are hereafter referred to as Treatment_(A+B)_ and Control_(A+B)_ expression. The results indicated that 1,155 (17.22%), 2,297 (32.56%), and 1,187 (17.45%) homoeologous pairs had Treatment_(A+B)_ expression values 2-fold greater than Control_(A+B)_ expression during three stress treatments (**Supplementary Table 68**), which were much less than the differentially expressed genes that were identified in traditional DEG analysis. The WGD increases gene redundancy and increases the expression plasticity of the allotetraploid genome, thus buffering gene expression in response to environmental stresses. The genes with significant expression changes could be buffered by the expression of its homoeologous copy and reduce the drastic impact of expression changes. Considering that the majority of homoeologous genes still maintain conserved expression patterns and gene functions, the DEG analysis based on summed expressions of homoeologous genes would provide an alternative approach for investigating gene regulations underlying phenotypic variation.

Taken together, the results of our expression divergence analysis using different methods provide substantial evidence of subgenome dominance and asymmetric expression, of which subgenome B not only retains more protein-coding genes but also has dominant gene expression levels under a relaxed purifying selection in the allotetraploid genome. However, it is still a challenging task to identify and discriminate among those mechanisms of gene functional divergence^42,43^, especially when accurate duplicate-aware gene annotation, well-powered transcriptomic analysis and expression studies, including embryonic stages, are absent for both allotetraploid *C. carpio* and the closely related diploids from its progenitor lineages.

### Methylation divergence in allopolyploid *C. carpio* genome

It is well recognized that DNA methylation is one of the important epigenetic regulation mechanisms for controlling gene expression^44^. To uncover DNA methylation divergence underlying expression divergence between two subgenomes, we generated single-base resolution methylomes from the YR strain of allotetraploid *C. carpio*. We identified 309,953,955 conserved cytosines with approximate 21,724,824 cytosines methylated (7.01%) present in three biological replicates for further analysis. We found that the cytosine methylation rates in subgenomes A and B were 6.88% and 7.13%, respectively, with no significant difference. The most abundant methylation occurs at CG sites in both subgenomes, with a CG methylation ratio of 85.98% in subgenome A and 86.34% in subgenome B, while the CHG and CHH methylation ratios are much lower (**Supplementary Table 69**). Therefore, we decided to focus on CG methylation in the methylation divergence analysis across two subgenomes. To test the relationship between methylation and sequence evolution in genic regions, we divided the 8,291 homoeologous gene pairs of two subgenomes into CG body-methylated (PCG < 0.05) and CG body-unmethylated (PCG > 0.95) genes using a binomial test with body-methylation levels. Among the 2,393 CG body-methylated homoeologous genes, the percentage of CG methylation changes was substantially higher than the substitution rate of the coding sequence (**Figure 4a**), which suggested that the methylation change rate is faster than the neutral sequence substitution rate. In the CG body-unmethylated genes, although the sequence variation remained at a similar level, the methylation peak disappeared. The faster methylation change rate suggested that, rather than vast gene loss and sequence substitution, epigenomic evolution provided quick and efficient regulation to deal with the massive changes of “genome shock” after the genome merger and ensured the survival of the tetraploidized *C. carpio*.

Many evolutionary studies on allotetraploid genomes suggested that dominantly expressed subgenomes tend to have less methylation sites in genic regions^45–48^. Therefore, we compared the global DNA methylation level of two subgenomes and looked for epigenetic evidence of subgenome dominance in *C. carpio* based on all annotated genes. We found that two subgenomes have similar CG methylation levels in the gene body and downstream regions, while subgenome A has a slightly higher CG methylation level in the upstream promotor regions than that of subgenome B (*p*<0.05) (**Figure 4b**). We also focused on the CG methylation of 8,291 homoeologous gene pairs to assess methylation divergence based on these highly-conserved genes. We found a similar result that the genes in subgenome A have a slightly higher CG methylation level in the promotor regions than that of their homoeologs in subgenome B (**Figure 4b**). The results suggested that the two subgenomes that were derived from different diploid progenitors retain a balanced CG methylation level in the gene body and downstream regions after the genome merger, which are not likely to be responsible for subgenome dominance and asymmetric expression in the allotetraploid genome. We reasoned that methylation changes in the promotor regions of the homoeologous gene pairs potentially regulate homoeologous expression bias and lead to subgenome dominance. Therefore, we further investigated CG methylation patterns among those that were divergently expressed homoeologous genes with an expression difference greater than 32-fold in 12 surveyed tissues. We found that the CG methylation levels in the promoter regions are much lower in the more highly expressed homoeologs than in the poorly expressed homoeologs in all 12 tissues, regardless of whether the higher expression homoeologs are in subgenome A or subgenome B. However, we did not observe a significant CG methylation difference in the gene body and downstream region between these asymmetrically expressed homoeologous gene pairs (**Supplementary Figure 18**). We further investigated the methylation divergence patterns of the homoeologous genes with extreme expression divergence, which are only expressed in one subgenome while they are silenced in the other (**Figure 4c**). The result indicated that the genes that were dominantly expressed in subgenome A retain a significantly lower level of CG methylation in the promoter region than their silenced homoeologs in subgenome B (*p* = 2.06e-07). Similarly, the dominantly expressed homoeologs in subgenome B also have a significantly reduced CG methylation level in the promoter region compared with their homoeologs in subgenome A (*p* = 4.23e-08). However, we did not find significant methylation divergence in the gene body or the downstream regions (**Figure 4d, Supplementary Figure 19**). As expected, the silenced *eif3i, acsl6* and *pd-l1*, as typical examples, have significantly increased CG methylation levels in the promoter regions than their normally expressed homoeologs (**Figure 4f**). Thus, we found a significant correlation between asymmetric expression and methylation divergence in the promoter region, which underpinned that CG methylation in the promoter regions plays an important role in altering the expression of these homoeologous genes in allotetraploid *C. carpio*. The selective methylation changes in homoeologous genes fine-tuned the gene expression to generate proper transcriptional products, which ensures the survival and evolutionary success of the tetraploidized *C. carpio*.

### Genetic basis of the reddish skin of domesticated Hebao red carp

Many studies have suggested that genome duplication plays a major role in shaping phenotypic diversity by increasing gene redundancy and shielding polyploids from the deleterious effect of mutations^26^. The common carp *C. carpio* has been recognized as one of the oldest domesticated fish in the world. Both natural and artificial selection have generated abundant phenotypic variations in both food and ornamental carp strains, such as the scale pattern, skin colour, body shape and adaptive plasticity to environmental stressors. It is a great challenge to investigate the genetic basis and regulate the mechanism of the phenotypic variations and adaptation in the complex tetraploid genome. Of the abundant variants of *C. carpio*, HB is one of the most renowned indigenous dual-purpose strains for food and ornamental uses in China due to its unique elliptical-shape body and reddish skin (**Supplementary Figure 1**). Hybridization experiments between the HB and wild type common carp suggested that skin colour was controlled by two independent genes^49^. Previous studies on the transcriptome expression analysis suggested that several candidate genes involving melanin synthesis were potentially associated with its reddish skin but led to controversy^13,50,51^. To unveil the genetic basis underlying the distinctive phenotypic variation of the skin colour of HB, we re-sequenced 34 random samples of domesticated HB and “wild-type” YR populations and identified 16.93 million SNPs between the two strains (**Supplementary Table 70**). The genome variation and allele frequency scans were subsequently performed and detected 248 (29.73 Mb) and 120 (11.08 Mb) genomic regions harbouring 1,237 and 470 candidate genes, with significant selection signatures found between the two strains (**Supplementary Table 71**). The gene, *bco2a-1*, was recognized from the candidate genes in chromosome B5 and encodes beta-carotene dioxygenase 2 (Bco2) (**Figure 4b**). Bco2 is an enzyme that is responsible for the asymmetric cleavage of β-carotene into β-apo-10’-carotenal and β-ionone and subsequently synthesizes vitamin A. Some studies previously confirmed that naturally occurring mutations in the *bco2* gene had a significant impact on the carotenoid metabolism and caused extra carotenoid deposition in various animals^52–54^. A 2-bp insertion was identified in exon 4 of the *bco2a-1* gene, exclusively in the HB population but not in the YR population (**Figure 4a**). We therefore reasoned that the 2-bp frameshift insertion in the coding region of *bco2a-1* is the potential causative mutation of the reddish skin of HB. Two *bco2* genes (*bco2a* and *bco2b)* were identified on chromosomes 5 and 15 in the diploid genome of zebrafish, which were likely derived from the Ts3R WGD. We identified four *bco2* genes (*bco2a-1, bco2a-2, bco2b-1*, and *bco2b-2*) in the allotetraploid *C. carpio* genome, of which homoeologous genes *bco2a-1* and *bco2a-2* on chromosome B5 and A5 are orthologues of *bco2a* of zebrafish, while homoeologous genes *bco2b-1* and *bco2b-2* on chromosomes B5 and B15 are orthologues of the *bco2b* of zebrafish. Likely, *bco2b-1* on chromosome B5 in subgenome B were translocated from chromosome A15 in subgenome A via homoeologous exchange after the recent genome merger in *C. carpio*. To determine whether asymmetrical expression also occurs in bco2 homoeologs, we measured the expression levels of all four *bco2* genes and found that *bco2a-1* is dominantly expressed, while all three of the other *bco2* genes were suppressed (**Figure 5b**), suggesting that the loss-of-function mutation of *bco2a-1* in the HB would substantially reduce the Bco2 level and interfere with carotenoid metabolism. Carotenoids accumulation may cause a reddish colour in the HB.

**Figure 5.**
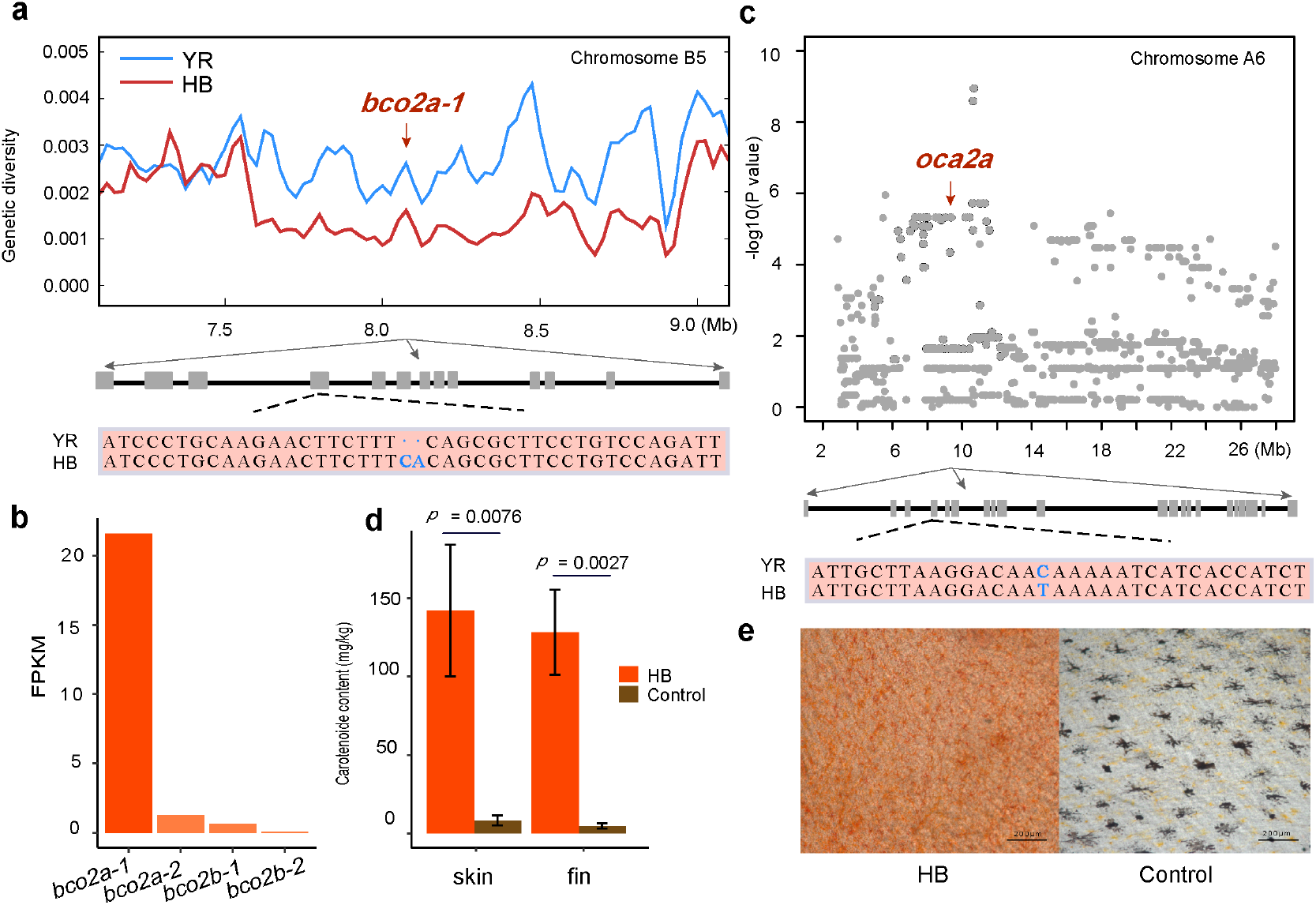
Genetic basis of skin colour phenotype in domesticated Hebao red carp. **a.** Highly divergent genomic region harbouring the *bco2a-1* gene in chromosome B5. The exons and introns of *bco2a-1* are represented by boxes and lines, respectively, and 2-bp insertion on the exon 2 is highlighted in blue. **b.** The relative expression levels of four *bco2* genes in the skin tissue of *C. carpio*, which indicate the dominant expression *bco2a-1* compared with its homoeologous and homologous copies. RPKM reads per kilobases per million, **c.** Local Manhattan plots obtained from the GWAS signals of the reddish skin colour trait in the introgression line from HB to YR. The gene *oca2a* is identified in the significant region. The missense C>T transversion mutation (T173I) in exon 4 is highlighted in blue. **d.** Histograms of carotenoid contents in reddish skin and fins of the HB and wild-type *C. carpio*. **f.** Melanocyte distribution in the skin of the HB and wild-type *C. carpio*.

To identify another potential genetic locus underlying the reddish skin of HB, we performed a genome-scale association study on an introgression line of the reddish skin phenotype that had been introgressed from HB to YR after continuous backcrosses at population level for 30 years. We found that approximately 1-2% individuals in the introgression line have reddish skin in contrast to the vast majorities with wild-type grey-brown skin (**Supplementary Figure 20**). We selected a family with a 3:1 phenotype segregation of wild-type and reddish skin from the introgression line for GWAS analysis, as was expected, and identified only one significant genome region on chromosome A06 (**Figure 5c, Supplementary Figure 21**). A candidate gene, *oca2a*, was identified in this region, which encodes the OCA2 melanosomal transmembrane protein (Oca2), which is essential for normal pigmentation and is likely to be involved in the production of melanin. Many previous studies had already indicated that mutations in the Oca2 gene cause a disruption in the normal production of melanin and therefore cause melanin reductions in the skin ^55–58^. A missense C>T transversion mutation was found in exon 4 of *oca2a*, which changes threonine to isoleucine (T173I) in Oca2. The missense mutation was presented as a homozygote only in the reddish YR from the introgression line but not in the wild-type YR population. We also found the homozygote mutation in HB population, which indicated that the point mutation was indeed derived from the HB population. The missense mutation is likely a loss-of-function mutation that caused skin melanin reduction in HB.

Thus, the carotenoid accumulation and/or melanin reduction caused by loss-of-function mutations would be the cause of reddish skin in the HB population. To test this hypothesis, we compared the carotenoid contents and melanocyte distribution in the reddish skin of HB with those of the grey-brown skin of regular common carp. We found that the carotenoid in the HB skin and fins was extraordinarily higher than that in control (**Figure 5d**). We also found that melanocytes were abundant in the grey-brown skin of regular common carp but were completely absent in the reddish skin of HB (**Figure 5e**). The observations support the hypothesis that the absence of melanin pigments allows accumulated carotenoids to display a bright red colour in the skin of HB.

## CONCLUSION

The allotetraploidized *C. carpio*, together with its close relatives with diverse ploidy in Cyprinidae, has been recognized as an excellent model system for studying polyploid genome evolution in vertebrates. Discovery of the possible subgenome origins for *C. carpio* will provide insight for further investigations of subgenome divergence and phenotypic adaptation of the newly formed allotetraploid teleost. In this study, we demonstrated substantial evidence and discovered a diploid lineage in Cyprinine as one of the potential progenitors of *C. carpio*. We therefore characterized the allotetraploid origin of *C. carpio* and divided its genome into two subgenomes marked by a distinct sequence similarity to that of the diploid progenitor. The subgenome merger was estimated to occur approximately 12.4 Mya in Southwest China during extensive uplift of the Qinghai-Tibetan Plateau. Asymmetric gene loss and rediploidization have been observed in many allopolyploid genomes. However, we found that the majority of homoeologous genes are well preserved in the *C. carpio* genome and did not observe large-scale gene loss and rediploidization. Instead, we discovered extensive homoeologous gene expression bias across twelve investigated tissues to show that subgenome B is more dominant in homoeologous expression than is subgenome A. Environmental stresses tend to increase homoeologous expression divergence. DNA methylation analysis suggested that CG methylation in promoter regions may play an important role in altering the expression of these homoeologous genes in allotetraploid *C. carpio*. The selective methylation changes in homoeologous genes fine-tuned gene expression to generate proper transcriptional products without massive gene losses and mutations, which ensured the survival and evolutionary success of the tetraploidized *C. carpio*. Our results also suggested that both the asymmetric expression of homoeologous genes and the missense mutations of *bco2a-1* and *oca2a* could be the genetic basis of the red skin colour phenotype in the domesticated Hebao red carp. This study provides insights for extending exploration on polyploidy evolution in vertebrates and will facilitate the improvement of economically important traits by focusing on dominantly expressed homoeologous genes and genome regions.

## METHODS

### Sample preparation, genome sequencing and assembly of common carp

We collected a female Yellow River carp (*C. carpio haematopterus*) at the Hatchery Station of Henan Academy of Fishery Sciences at Zhengzhou; a female Hebao red carp (*C. carpio wuyuanensis*) at Wuyuan County in Jiangxi Province, China; and a female German Mirror carp (*C. carpio carpio*) at Heilongjiang Fishery Research Institute in Harbin as genomic DNA donors for whole genome sequencing. Genomic DNA was extracted from blood using a QIAGEN DNeasy Blood & Tissue Kit (QIAGEN, Shanghai, China). We constructed eight sequencing libraries for three carps with various insert sizes from 250 bp to 20 Kbp according to Illumina standard operating procedures. Subsequently, we used the Illumina HiSeq platform to sequence these libraries with 150-bp read length, and generated 339.11 Gb, 298.65 Gb, and 330.39 Gb clean data for three genome assemblies. We estimated the genome size based on 17-mer frequency distribution and assembled the genome via a modified version of SOAP*denovo*, specifically for the high heterozygous genome^59^. To evaluate the accuracy of the assemblies at single base level, we mapped short sequence reads to carp genomes with BWA^60^ and performed variant calling with SAMtools^61^. We also assessed assembly completeness by remapping the PE reads to CEGMA and ESTs. To assess the genome connectivity and assembly accuracy, we aligned 34,932 previously published mate-paired BAC-end sequences (BES) derived from Songpu mirror carp to the new assemblies^62,63^.

We constructed high-density genetic linkage maps for HB and GM by genotyping F1 mapping families on the high-density 250K SNP array following Affymetrix protocols. The double pseudo-test cross strategy was employed for linkage analysis using JoinMap 4.1 (https://www.kyazma.nl/). Recombination frequencies of markers on the same LG were converted into map distances (cM) through the maximum likelihood (ML) algorithm. The consensus map was then established using the MergeMap by integrating sex-specific maps through shared markers. All genetic linkage maps were drawn using MapChart 2.2^64^. Together with previously published high-density genetic linkage maps of YR, we have three high-density linkage maps available for scaffold integration.

To anchor scaffolds to each linkage map, we aligned the SNP-associated sequences of high-density genetic linkage maps to the assembled genomes using BLAST. Only SNP markers with a unique location were used for anchoring and orienting scaffolds. For those scaffolds that were in conflict with the genetic map, we performed manual checks using mate-paired reads.

### Gene prediction and functional annotation

Gene prediction and functional annotation were performed through a combination of homology-based prediction, *de novo* prediction and transcriptome-based prediction methods. Protein sequences from *Ctenopharyngodon idellus, Cynoglossus semilaevis, Denio rerio, Oryzias latipes, Tetraodon nigroviridis, Sinocyclocheilus graham, Sinocyclocheilus rhinocerous, Sinocyclocheilus anshuiensis, Mus musculus* and *Homo sapiens* were aligned to the carp genome using TblastN (E-value <= le-5). The BLAST hits were conjoined by GeneWise (v2.4.1)^65^ for accurate spliced alignments. For *de novo* prediction, three *de novo* prediction tools, Augustus (v2.7)^66^, GlimmerHMM (v3.02)^67^ and SNAP (version 2006-07-28)^68^, were used to predict the genes in the repeat-masked genome sequences. The RNA-seq reads from multiple tissues were mapped onto the genome assembly using Tophat (v2.1.0)^69^, and then Cufflinks (v2.1.1)^70^ was used to assemble the transcripts into gene models. Gene predictions from the *de novo* approach, *homology-based* approach and RNA-Seq-based evidence were merged to form a comprehensive consensus gene set using the software EVM^71^. To achieve the functional annotation, the predicted protein sequences were aligned against public databases, including SwissProt, TrEMBLE and KEGG with BLASTP (E-value<=le-5). Additionally, protein motifs and domains were annotated by searching the InterPro and Gene Ontology (GO) databases using InterProScan (v4.8)^72^.

### Repetitive element annotation

Transposable elements in carp genomes were detected by combining homology-based and *de novo* predictions. For homology-based detection, RepeatMasker and RepeatProteinMask were used to screen the carp genome for known transposable elements against the RepBase library (v20140131) (http://www.repeatmasker.org/). *De novo* transposable elements in the genome were identified by RepeatMasker based on a *de novo* repeat library constructed by RepeatModeller and LTR_FINDER (v1.0.5)^73^. Tandem repeats were detected using the program Tandem Repeats Finder (TRF, v4.07b)^74^ with default parameters.

### RNA sequencing

Twelve tissues (brain, muscle, liver, intestine, blood, head kidney, trunk kidney, skin, gill, spleen, gonad and heart) were dissected and collected from six Yellow River carp. Total RNA was extracted from 12 tissues using TRIZOL (Invitrogen, Carlsbad, CA, USA). The RNA samples were then treated by DNase I. The integrity and size distribution were checked with Bioanalyzer 2100 (Agilent technologies, Santa Clara, CA, USA). The high-quality RNA samples were sequenced on Illumina HiSeq 2000 platforms with the manufacturer’s instructions. A total of 72.2 Gb clean reads were generated for 12 tissues for expression analysis. Similarly, we also collected 75.6 Gb RNA reads of the same 12 tissues from grass carp *C. idellus*, which has the complete reference genome available and serves as a diploid reference in this study. The RNA-seq data for biotic and abiotic stresses treatment were collected from SRA databases (Accession No. PRJNA314552 for CyHV-3 infection, Accession No. PRJNA315069 for *Aeromonas hydrophila* infection, and No. PRJNA512071 for hypoxia experiment).

### Phylogenetic analysis

To explore the evolutionary relationship of *C. carpio* and its closely related tetraploid and diploid Cyprininae species, we constructed an ML phylogenetic tree based on Rag2 genes selected from a previous published dataset^12^ or extracted from published Cyprinid genomes in software MEGA7^75^. Four diploid Cyprininae species *(Poropuntius huangchuchieni, Hampala macrolepidota, Onychostoma barbatulum*, and *Cirrhinus molitorella*) representing three closely related diploid clades were selected for further investigation.

To confirm the homoeologous relationship of the two subgenomes in *C. carpio*, we built phylogenetic trees based on 2,071 conserved homoeologous gene pairs among two tetraploids (*C. carpio* and *S. anshuiensis*) and their orthologues from three diploids *D. rerio, C. idella* and *P. huangchuchieni*) using RAxML software^76^ and integrated 2,071 trees using programs MP-EST^77^ and summarytree based on their topologies.

### Sample collection and genome sequencing of four Cyprinid species

Four diploid Cyprinid species that are closely related to allotetraploid common carp, *P. huangchuchieni, H. macrolepidota, O. barbatulum*, and *C. molitorella*, were collected for whole genome sequencing and draft assembly. *P. huangchuchieni* and *H. macrolepidota* were collected at Xishuangbanna in Yunnan Province, China, *Onychostoma barbatulum* was collected at Lishui in Zhejiang Province, China, and *Cirrhinus molitorella* was collected at Guangzhou, Guangdong Province, China. Genomic DNA was extracted from a fin clip or muscle using QIAGEN DNeasy Blood & Tissue Kit. The 350-bp whole-genome libraries were constructed for each sample according to the manufacturer’s specifications (Illumina). The libraries were then sequenced on the Illumina HiSeq platform to generate raw sequences with a 150-bp read length. A total of 54.68 Gb (*Cirrhinus molititorella*), 57.35 Gb (*Poropuntius huangchuchieni*), 64.31 Gb (*Onychostoma barbatula*) and 71.78 Gb (*Hampala macrolepidota*) data were generated for the above four species and were subjected for primary genome assemblies using SOAP*denovo*.

### Defining the conserved homoeologous gene pairs

To determine the conserved homoeologous gene pairs in the tetraploid genome of *C. carpio*, we used annotated protein coding genes of the *C. idella* genome as a diploid reference. The OrthoMCL pipeline^78^ was used to define gene families in the common ancestor. The all-against-all similarities were determined using blastp with an E-value cutoff of 1e-5. The orthologous triplets with 1:1:1 relationship in *C. idella* genome, subgenomes A and B of *C. carpio* were identified from two genomes. The orthologous gene pairs in the *C. carpio* genome were then defined as homoeologous genes that were derived from the latest WGD event.

### Homoeologous gene expression analysis

Clean reads of RNA-seq were mapped onto the reference genome of *C. carpio* using Hisat2 (v2.0.4)^79^. Gene expressed level in terms of reads per kilobase of transcript per million mapped reads (RPKM) was estimated by HTseq^80^ and custom Perl scripts. Samples and genes were clustered using Pearson’s correlation and Ward’s method in the R function hclust and were visualized as heatmaps using the R function ggplot2.

A homoeolog expression dominance analysis was performed for syntenic gene pairs. Differentially expressed genes pairs with greater than twofold change were defined as dominant gene pairs. The dominant genes were the genes that were expressed relatively higher in dominant gene pairs, and the lower ones were the subordinate genes. The rest of the syntenic gene pairs that showed non-dominance were classified as neutral genes.

### Whole genome bisulfite sequencing (WGBS)

Genomic DNA was extracted from muscle tissue of three common carp individuals. Genomic DNA was treated using a sodium bisulfite, which converts unmethylated cytosine to uracil, then thymine^81^. Whole-genome bisulfite sequencing (WGBS) was performed using an Illumina HiSeq2500 sequencer with 150 bp paired-end sequences (Illumina, San Diego, CA, USA).

### Whole-genome resequencing, SNP calling, and genome diversity screening

Sixteen samples of YR and eighteen samples of HB, respectively, were collected randomly and were sequenced using the Illumina HiSeq 2000 platform. Paired end reads from each sample were aligned to the HH reference genome using the Burrows-Wheeler Aligner (BWA). SNPs were identified from the VCF files that were generated with SAMtools. The filtering threshold was set to read at a depth ≥10 and a quality score ≥20. All SNPs were used to investigate the population structure using STRUCTURE^82^ with 2,000 iterations. The result of the structure matrix was plotted using STRUCTURE PLOT V2.0^83^ software. Linkage disequilibrium was calculated using PLINK^84^ with an R^2^ threshold of >0.2, and LD decay was calculated by R2 values against distances among the SNPs.

We calculated the Pi distribution and Wright’s fixation index (FST) for each linkage group using a sliding window method with VCFTOOLS software^85^. The window width was set to 10 kb, and the stepwise distance was 1 kb. Furthermore, to identify selection signatures more accurately, we used the extended haplotype homozygosity (XP-EHH) statistic to infer the selected regions compared to a reference population.

### GWAS

We performed a GWAS using GEMMA (Genome-wide Efficient Mixed Model Association)^86^ with univariate linear models. The Manhattan plot of the −log 10 (P value) was accomplished by an R package. The genome-wide significance for GWAS was defined according to the Bonferroni method as 0.05/N, where N was the total number of SNPs used in our study^87^.

### Carotenoids assessment and melanocyte observation

Skin and fin tissues were collected from Hebao red carp and the wild-type control. The tissues were rinsed in 0.7% saline solution and then were observed for examining chromatophore distribution on a Zeiss Axio imager A2 microscope. Carotenoid contents were measured at an absorbance of 476 nm using a PerkinElmer Lambda 950 UV-vis spectrometer according to previous published method^88^.

## Acknowledgements

We acknowledge grant support from the National Natural Science Foundation of China (grants 31422057 & 31872561 to P.X. and 31502151 to J.X.), the National High-Technology Research and Development Program of China (grant 2011AA100401 to P.X.), the Fundamental Research Funds for the Central Universities, Xiamen University (grants. 20720180123 & 20720160110 to P.X.), Central Public-interest Scientific Institution Basal Research Fund, CAFS (No.2016HY-JC0301 to J.X.) and the National Key Research and Development Program (2018YFD0900102 to J.X.).

## Contributions

P.X. conceived and designed the research. P.X. and J.X. coordinated the project. G.L. and W. J. developed the sequencing strategy, performed next-generation sequencing, assembly and bioinformatics. J.X., L.C. Z. Zhao and C.D. performed RNA-Seq collection and DNA methylation sequencing. P.X., J.X., L.C., Z. Zhou, Y.J., Z. Zhao, B.C., F.P., J.F., J.L., H.W. and C.D. collected and prepared the samples. J.X., L.C., W.P, and Z.J. performed the genotyping and genetic linkage mapping. Z. Zhou, W.P., and L.C. performed phylogenetic analysis. Z. Zhou and Y.J. performed GWAS analysis. J.X., B.C. and H.B. performed wet-lab molecular experiments. Y.W. and Y.S. validated skin color phenotype. P.X., L.C., G.L. and J.X. wrote and revised the manuscript and the supplementary information. X.S. participated in discussions and provided valuable advice.

## Data availability

All the genomic sequence datasets have been deposited in the NCBI Sequence Read Archive under accession number PRJNA510861, PRJNA511029-PRJNA511032.

## References

1. David, L., Blum, S., Feldman, M.W., Lavi, U. & Hillel, J. Recent duplication of the common carp (Cyprinus carpio L.) genome as revealed by analyses of microsatellite loci. Mol Biol Evol 20, 1425–34 (2003).

2. Larhammar, D. & Risinger, C. Molecular genetic aspects of tetraploidy in the common carp Cyprinus carpio. Mol Phylogenet Evol 3, 59–68 (1994).

3. Postlethwait, J.H. et al. Zebrafish comparative genomics and the origins of vertebrate chromosomes. Genome Res 10, 1890–902 (2000).

4. Van de Peer, Y., Taylor, J.S., Braasch, I. & Meyer, A. The ghost of selection past: rates of evolution and functional divergence of anciently duplicated genes. J Mol Evol 53, 436–46 (2001).

5. Vandepoele, K., De Vos, W., Taylor, J.S., Meyer, A. & Van de Peer, Y. Major events in the genome evolution of vertebrates: paranome age and size differ considerably between ray-finned fishes and land vertebrates. Proceedings of the National Academy of Sciences of the United States of America 101, 1638–1643 (2004).

6. Ohno, S. Evolution by gene duplication, xv, 160 p. (Springer-Verlag, Berlin, New York, 1970).

7. Allendorf, F.W. & Thorgaard, G.H. Tetraploidy and the evolution of salmonid fishes, in Evolutionary genetics of fishes 1–53 (Springer, 1984).

8. Phillips, R. & Rab, P. Chromosome evolution in the Salmonidae (Pisces): an update. Biological Reviews of the Cambridge Philosophical Society 76, 1–25 (2001).

9. Ohno, S., Muramoto, J., Christian, L. & Atkin, N.B. Diploid-tetraploid relationship among old-world members of the fish family Cyprinidae. Chromosoma 23, 1–9 (1967).

10. Wolf, U., Ritter, H., Atkin, N. & Ohno, S. Polyploidization in the fish family Cyprinidae, order Cypriniformes. Humangenetik 7, 240–244 (1969).

11. Kumar, S., Stecher, G., Suleski, M., Hedges, S.B.J.M.B. & Evolution. TimeTree: a resource for timelines, timetrees, and divergence times. 34, 1812–1819 (2017).

12. Wang, X., Gan, X., Li, J., Chen, Y. & He, S. Cyprininae phylogeny revealed independent origins of the Tibetan Plateau endemic polyploid cyprinids and their diversifications related to the Neogene uplift of the plateau. Science China Life Sciences 59, 1149–1165 (2016).

13. Xu, P. et al. Genome sequence and genetic diversity of the common carp, Cyprinus carpio. Nat Genet 46, 1212–9 (2014).

14. Liu, S. et al. Genomic incompatibilities in the diploid and tetraploid offspring of the goldfish x common carp cross. Proc Natl Acad Sci USA 113, 1327–32 (2016).

15. Chen, Z. et al. De Novo assembly of the goldfish (<em>Carassius auratus</em>) genome and the evolution of genes after whole genome duplication. (2018).

16. Bostock, J. et al. Aquaculture: global status and trends. Philosophical Transactions of the Royal Society B: Biological Sciences 365, 2897–2912 (2010).

17. Fisheries, F. Aquaculture Department (2007) The State of World Fisheries and Aquaculture 2006. Food and Agriculture Organization of the United Nations, Rome.

18. Xu, J. et al. Development and evaluation of the first high-throughput SNP array for common carp (Cyprinus carpio). BMC Genomics 15, 307 (2014).

19. Nelson, J.S. Fishes of the World. 4th. Hoboken: Wiley (2006).

20. Howes, G. Systematics and biogeography: an overview, in Cyprinid fishes 1–33 (Springer, 1991).

21. Zhao, L. et al. A dense genetic linkage map for common carp and its integration with a BAC-based physical map. PLoS One 8, e63928 (2013).

22. Peng, W. et al. An ultra-high density linkage map and QTL mapping for sex and growth-related traits of common carp (Cyprinus carpio). Sci Rep 6, 26693 (2016).

23. Willett, C.E., Cherry, J.J. & Steiner, L.A. Characterization and expression of the recombination activating genes (ragl and rag2) of zebrafish. Immunogenetics 45, 394–404 (1997).

24. Favre, A. et al. The role of the uplift of the Qinghai-Tibetan Plateau for the evolution of Tibetan biotas. Biological Reviews 90, 236–253 (2015).

25. Madlung, A. Polyploidy and its effect on evolutionary success: old questions revisited with new tools. Heredity (Edinb) 110, 99–104 (2013).

26. Comai, L. The advantages and disadvantages of being polyploid. Nature Reviews Genetics 6, 836–846 (2005).

27. Edger, P.P., McKain, M.R., Bird, K.A. & VanBuren, R. Subgenome assignment in allopolyploids: challenges and future directions. Curr Opin Plant Biol 42, 76–80 (2018).

28. Cheng, F. et al. Gene retention, fractionation and subgenome differences in polyploid plants. Nat Plants A, 258–268 (2018).

29. Edger, P.P. & Pires, J.C. Gene and genome duplications: the impact of dosage-sensitivity on the fate of nuclear genes. Chromosome Research 17, 699 (2009).

30. Jiao, Y. & Paterson, A.H. Polyploidy-associated genome modifications during land plant evolution. Phil. Trans. R. Soc. B 369, 20130355 (2014).

31. Blanc, G. & Wolfe, K.H. Functional divergence of duplicated genes formed by polyploidy during Arabidopsis evolution. The Plant Cell 16, 1679–1691 (2004).

32. Aravind, L., Watanabe, H., Lipman, D.J. & Koonin, E.V. Lineage-specific loss and divergence of functionally linked genes in eukaryotes. Proceedings of the National Academy of Sciences 97, 11319–11324 (2000).

33. Wang, M. et al. Asymmetric subgenome selection and cis-regulatory divergence during cotton domestication. Nat Genet 49, 579–587 (2017).

34. Lien, S. et al. The Atlantic salmon genome provides insights into rediploidization. Nature 533, 200-+ (2016).

35. Session, A.M. et al. Genome evolution in the allotetraploid frog Xenopus laevis. Nature 538, 336–343 (2016).

36. Pophaly, S.D. & Tellier, A. Population Level Purifying Selection and Gene Expression Shape Subgenome Evolution in Maize. Mol Biol Evol 32, 3226–35 (2015).

37. Aitken, C.E. & Lorsch, J.R. A mechanistic overview of translation initiation in eukaryotes. Nat Struct Mol Biol 19, 568–76 (2012).

38. Gomes-Duarte, A., Lacerda, R., Menezes, J. & Romao, L. elF3: a factor for human health and disease. RNA Biol 15, 26–34 (2018).

39. Chen, L. & Han, X. Anti-PD-1/PD-Ll therapy of human cancer: past, present, and future. J Clin Invest 125, 3384–91 (2015).

40. Miller, R.A., Miller, T.N. & Cagle, P.T. PD-1/PD-L1, Only a Piece of the Puzzle. Arch Pathol Lab Med (2016).

41. Teodoro, B.G. et al. Long-chain acyl-CoA synthetase 6 regulates lipid synthesis and mitochondrial oxidative capacity in human and rat skeletal muscle. J Physiol 595, 677–693 (2017).

42. Braasch, I., Bobe, J., Guiguen, Y. & Postlethwait, J.H. Reply to: ‘Subfunctionalization versus neofunctionalization after whole-genome duplication’. Nature genetics 50, 910 (2018).

43. Sandve, S.R., Rohlfs, R.V. & Hvidsten, T.R. Subfunctionalization versus neofunctionalization after whole-genome duplication. Nature genetics 50, 908 (2018).

44. Phillips, T. The role of methylation in gene expression. Nature Education 1, 116 (2008).

45. Song, Q., Zhang, T., Stelly, D.M. & Chen, Z.J. Epigenomic and functional analyses reveal roles of epialleles in the loss of photoperiod sensitivity during domestication of allotetraploid cottons. Genome Biol 18, 99 (2017).

46. Wang, X. et al. Gene-body CG methylation and divergent expression of duplicate genes in rice. Sci Rep 7, 2675 (2017).

47. Jackson, S.A. Epigenomics: dissecting hybridization and polyploidization. Genome Biol 18, 117 (2017).

48. Jiao, W. et al. Asymmetrical changes of gene expression, small RNAs and chromatin in two resynthesized wheat allotetraploids. Plant J 93, 828–842 (2018).

49. Zhang, J. & Pan, G. Body form and body colour in hybrids of Cyprinus carpio. Journal of Fisheries of China 7, 301–312 (1983).

50. Jiang, Y. et al. Comparative transcriptome analysis reveals the genetic basis of skin color variation in common carp. PloS one 9, e108200 (2014).

51. Kottler, V.A., Künstner, A. & Schartl, M. Pheomelanin in fish? Pigment cell & melanoma research 28, 355–356 (2015).

52. Berry, S.D. et al. Mutation in bovine beta-carotene oxygenase 2 affects milk color. Genetics 182, 923–6 (2009).

53. Vage, D.l. & Boman, I.A. A nonsense mutation in the beta-carotene oxygenase 2 (BCO2) gene is tightly associated with accumulation of carotenoids in adipose tissue in sheep (Ovis aries). Bmc Genetics 11(2010).

54. Toews, D.P.L., Hofmeister, N.R. & Taylor, S.A. The Evolution and Genetics of Carotenoid Processing in Animals. Trends in Genetics 33, 171–182 (2017).

55. Klaassen, H., Wang, Y., Adamski, K., Rohner, N. & Kowalko, J.E. CRISPR mutagenesis confirms the role of oca2 in melanin pigmentation in Astyanax mexicanus. Dev Biol (2018).

56. Saenko, S.V. et al. Amelanism in the corn snake is associated with the insertion of an LTR-retrotransposon in the OCA2 gene. Scientific Reports 5(2015).

57. Protas, M.E. et al. Genetic analysis of cavefish reveals molecular convergence in the evolution of albinism. Nature Genetics 38, 107–111 (2006).

58. Fukamachi, S. et al. Conserved function of medaka pink-eyed dilution in melanin synthesis and its divergent transcriptional regulation in gonads among vertebrates. Genetics 168, 1519–1527 (2004).

59. Li, R.Q. et al. De novo assembly of human genomes with massively parallel short read sequencing. Genome Research 20, 265–272 (2010).

60. Li, H. & Durbin, R. Fast and accurate short read alignment with Burrows-Wheeler transform. Bioinformatics 25, 1754–60 (2009).

61. Li, H. et al. The Sequence Alignment/Map format and SAMtools. Bioinformatics 25, 2078–9 (2009).

62. Xu, P. et al. Genomic insight into the common carp (Cyprinus carpio) genome by sequencing analysis of BAC-end sequences. Bmc Genomics 12(2011).

63. Li, Y. et al. Construction and Characterization of the BAC Library for Common Carp Cyprinus Carpio L. And Establishment of Microsynteny with Zebrafish Danio Rerio. Marine Biotechnology 13, 706–712 (2011).

64. Voorrips, R. MapChart: software for the graphical presentation of linkage maps and QTLs. Journal of heredity 93, 77–78 (2002).

65. Birney, E. & Durbin, R. Using GeneWise in the Drosophila annotation experiment. Genome Research 10, 547–548 (2000).

66. Stanke, M., Schöffmann, O., Morgenstern, B. & Waack, S. Gene prediction in eukaryotes with a generalized hidden Markov model that uses hints from external sources. BMC bioinformatics 7, 62 (2006).

67. Salzberg, S.L., Delcher, A.L., Kasif, S. & White, O. Microbial gene identification using interpolated Markov models. Nucleic acids research 26, 544–548 (1998).

68. Korf, I.B.b. Gene finding in novel genomes. BMC bioinformatics 5, 59 (2004).

69. Kim, D. etal. TopHat2: accurate alignment of transcriptomes in the presence of insertions, deletions and gene fusions. Genome biology 14, R36 (2013).

70. Trapnell, C. etal. Transcript assembly and quantification by RNA-Seq reveals unannotated transcripts and isoform switching during cell differentiation. Nature biotechnology 28, 511 (2010).

71. Haas, B.J. et al. Automated eukaryotic gene structure annotation using EVidenceModeler and the Program to Assemble Spliced Alignments. Genome biology 9, 1 (2008).

72. Zdobnov, E.M. & Apweiler, R. InterProScan-an integration platform for the signature-recognition methods in InterPro. Bioinformatics 17, 847–848 (2001).

73. Xu, Z. & Wang, H. LTR_FINDER: an efficient tool for the prediction of full-length LTR retrotransposons. Nucleic acids research 35, W265–W268 (2007).

74. Benson, G. Tandem repeats finder: a program to analyze DNA sequences. Nucleic acids research 27, 573–580 (1999).

75. Kumar, S., Stecher, G., Tamura, K. & evolution. MEGA7: molecular evolutionary genetics analysis version 7.0 for bigger datasets. Molecular biology 33, 1870–1874 (2016).

76. Stamatakis, A. RAxML version 8: a tool for phylogenetic analysis and post-analysis of large phylogenies. Bioinformatics 30, 1312–1313 (2014).

77. Liu, L., Yu, L. & Edwards, S.V. A maximum pseudo-likelihood approach for estimating species trees under the coalescent model. BMC evolutionary biologyV 10, 302 (2010).

78. Li, L., Stoeckert, C.J. & Roos, D.S. OrthoMCL: identification of ortholog groups for eukaryotic genomes. Genome research 13, 2178–2189 (2003).

79. Kim, D., Langmead, B. & Salzberg, S.L. HISAT: a fast spliced aligner with low memory requirements. Nat Methods 12, 357–60 (2015).

80. Anders, S., Pyl, P.T. & Huber, W. HTSeq--a Python framework to work with high-throughput sequencing data. Bioinformatics 31, 166–9 (2015).

81. Wang, Y., Zheng, W.L., Luo, J.F., Zhang, D.D. & Lu, Z.H. In situ bisulfite modification of membrane-immobilized DNA for multiple methylation analysis. Analytical Biochemistry 359, 183–188 (2006).

82. Pritchard, J.K., Stephens, M. & Donnelly, P. Inference of population structure using multilocus genotype data. Genetics 155, 945–959 (2000).

83. Ramasamy, R.K., Ramasamy, S., Bindroo, B.B. & Naik, V.G. STRUCTURE PLOT: a program for drawing elegant STRUCTURE bar plots in user friendly interface. SpringerPlus 3, 431 (2014).

84. Purcell, S. etal. PLINK: A Tool Set for Whole-Genome Association and Population-Based Linkage Analyses. The American Journal of Human Genetics 81, 559–575 (2007).

85. Danecek, P. et al. The variant call format and VCFtools. Bioinformatics 27, 2156–2158 (2011).

86. Zhou, X. & Stephens, M. Genome-wide efficient mixed-model analysis for association studies. Nature Genetics 44, 821 (2012).

87. Gutierrez, A.P., Yáñez, J.M., Fukui, S., Swift, B. & Davidson, W.S. Genome-Wide Association Study (GWAS) for Growth Rate and Age at Sexual Maturation in Atlantic Salmon (Salmo salar). Plos One 10, e0119730 (2015).

88. Jiang, Z.-q. et al. Deposition and distribution of carotenoids in different tissues of ornamental carp Cyprinus carpio [J], Journal of Dalian Ocean University 1, 007 (2012).

