## Supplementary File 1 for "The allotetraploid origin and asymmetrical genome evolution of common carp *Cyprinus carpio*"

### SUPPLEMENTARY INFORMATION

### Supplementary Figures

**Supplementary Figure 1. The photos of three sequenced common carp strains.**

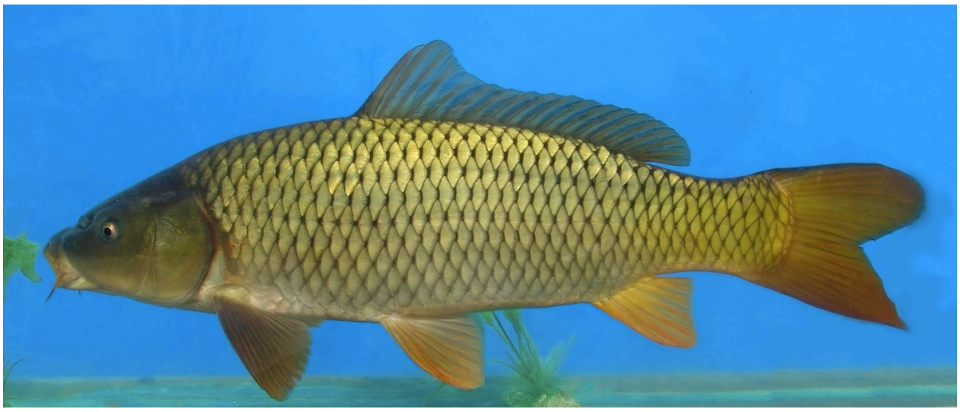

**(A) Yellow River Carp**

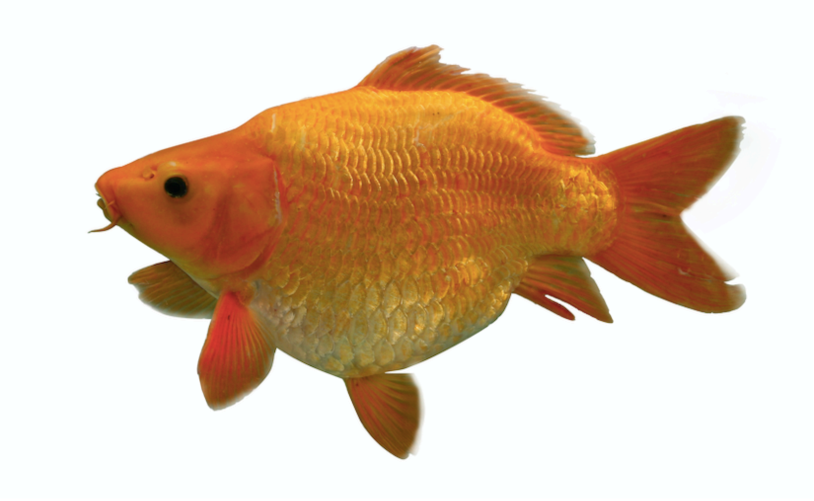

**(B) Hebao Red Carp**

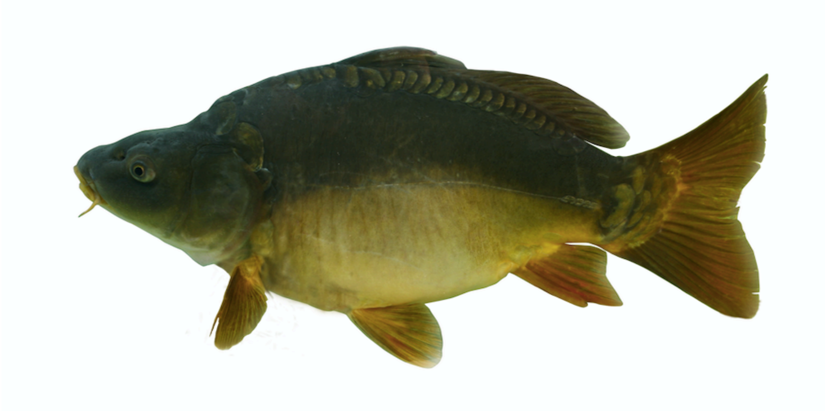

**(C) German Mirror Carp**

**Supplementary Figure 2. High density genetic linkage maps of three subspecies of *C. carpio*.**

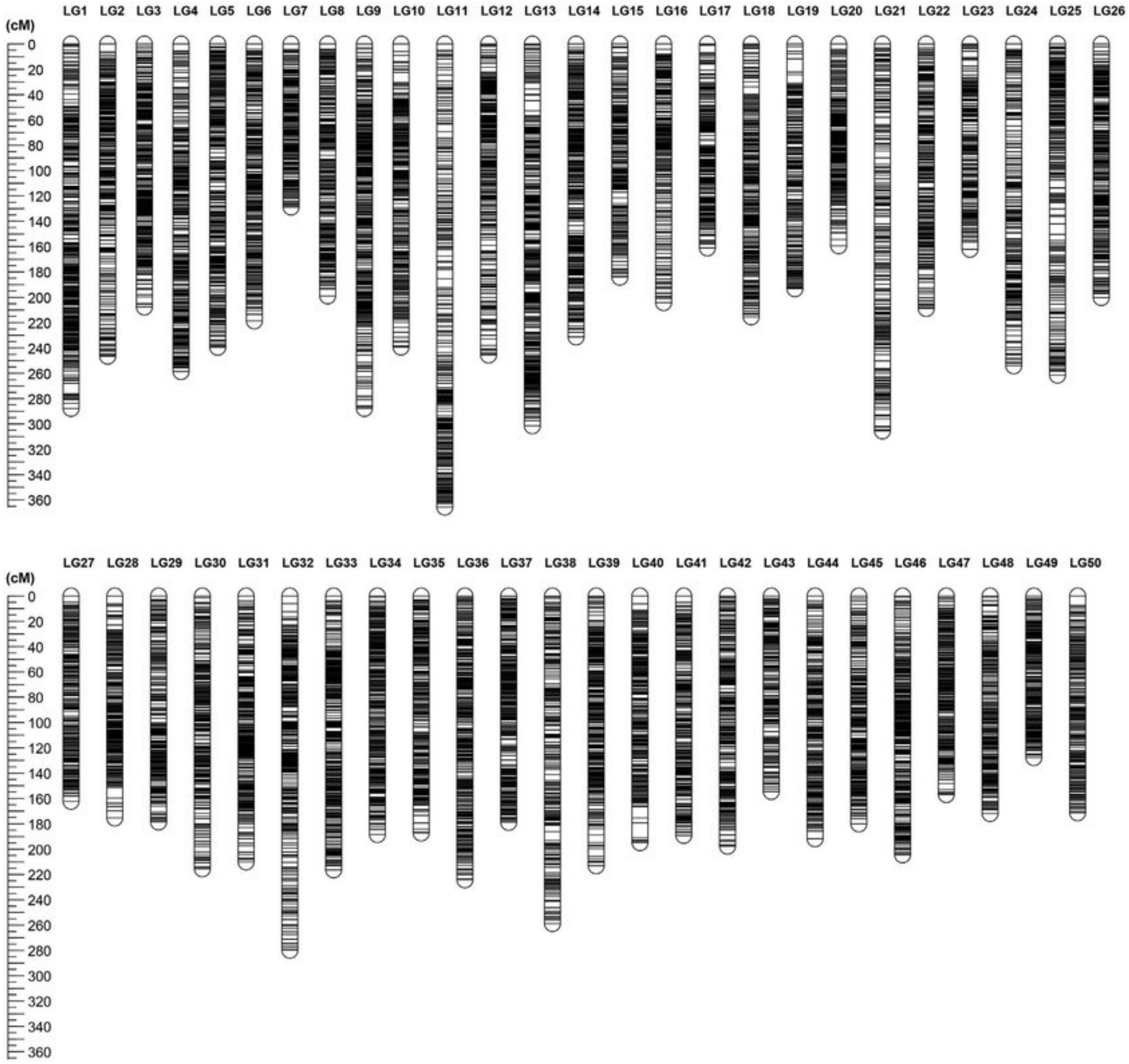

(A) Genetic linkage map of Yellow River carp

Note: The genetic linkage map for Yellow River carp had been previously published on Scientific Reports (Peng, et al, 2016)

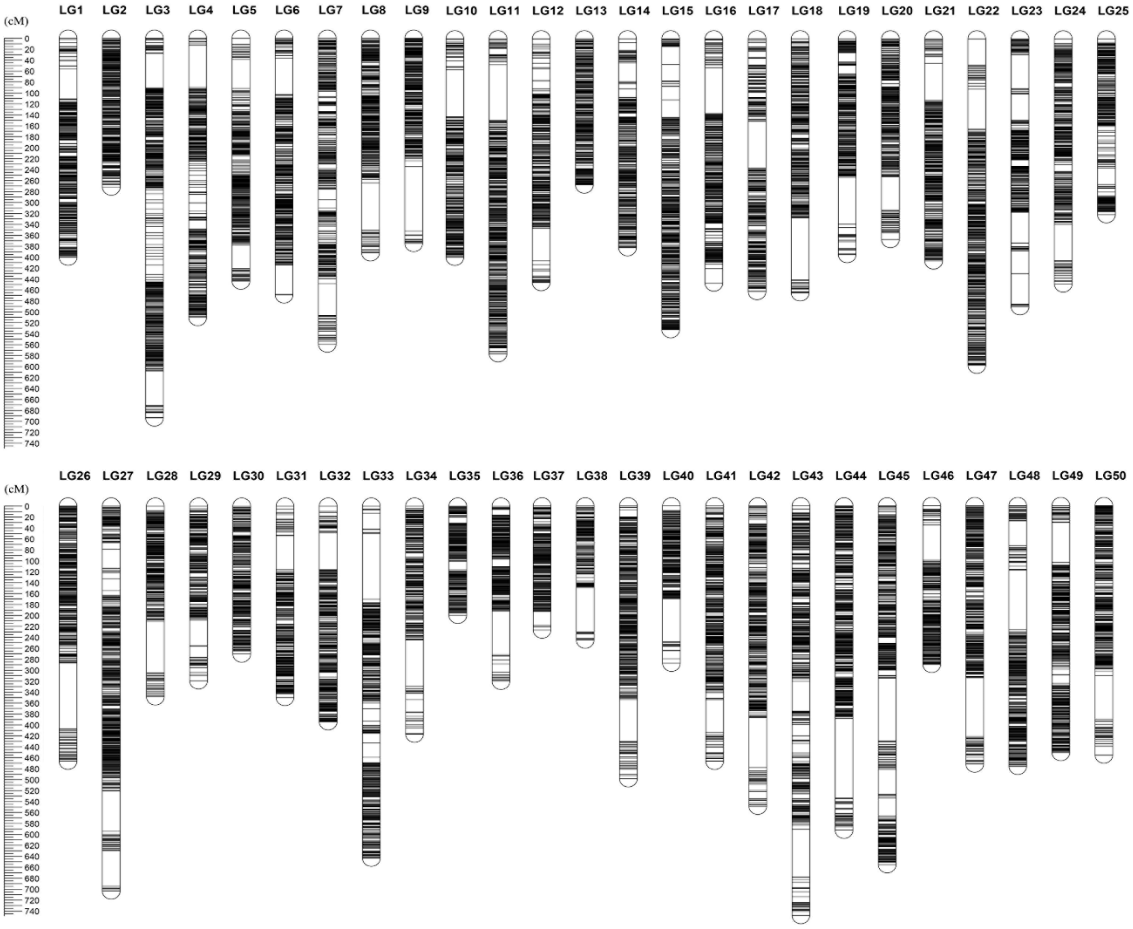

(B) Genetic linkage map of Hebao red carp

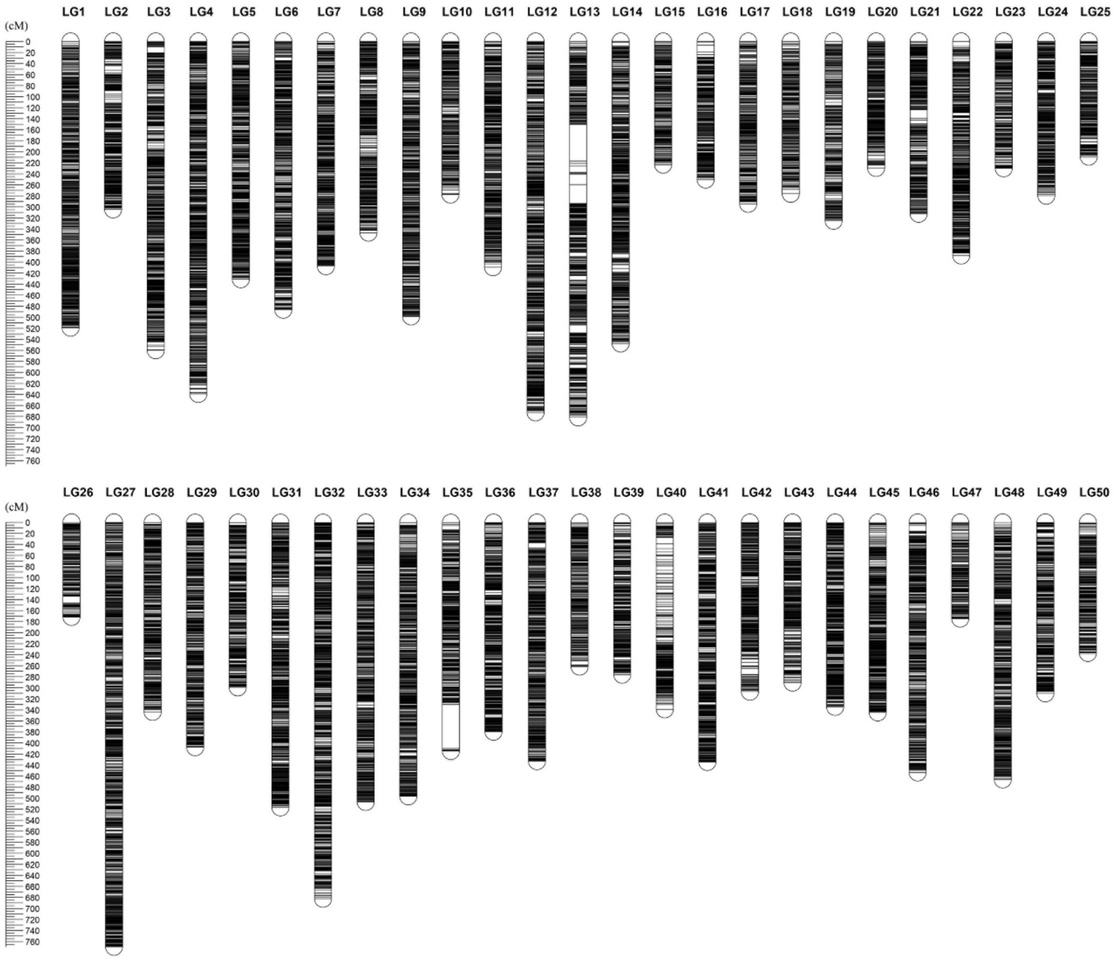

(C) Genetic linkage map of German mirror carp

**Supplementary Figure 3. Comparison between GM and SP Based on BAC-end sequences.**

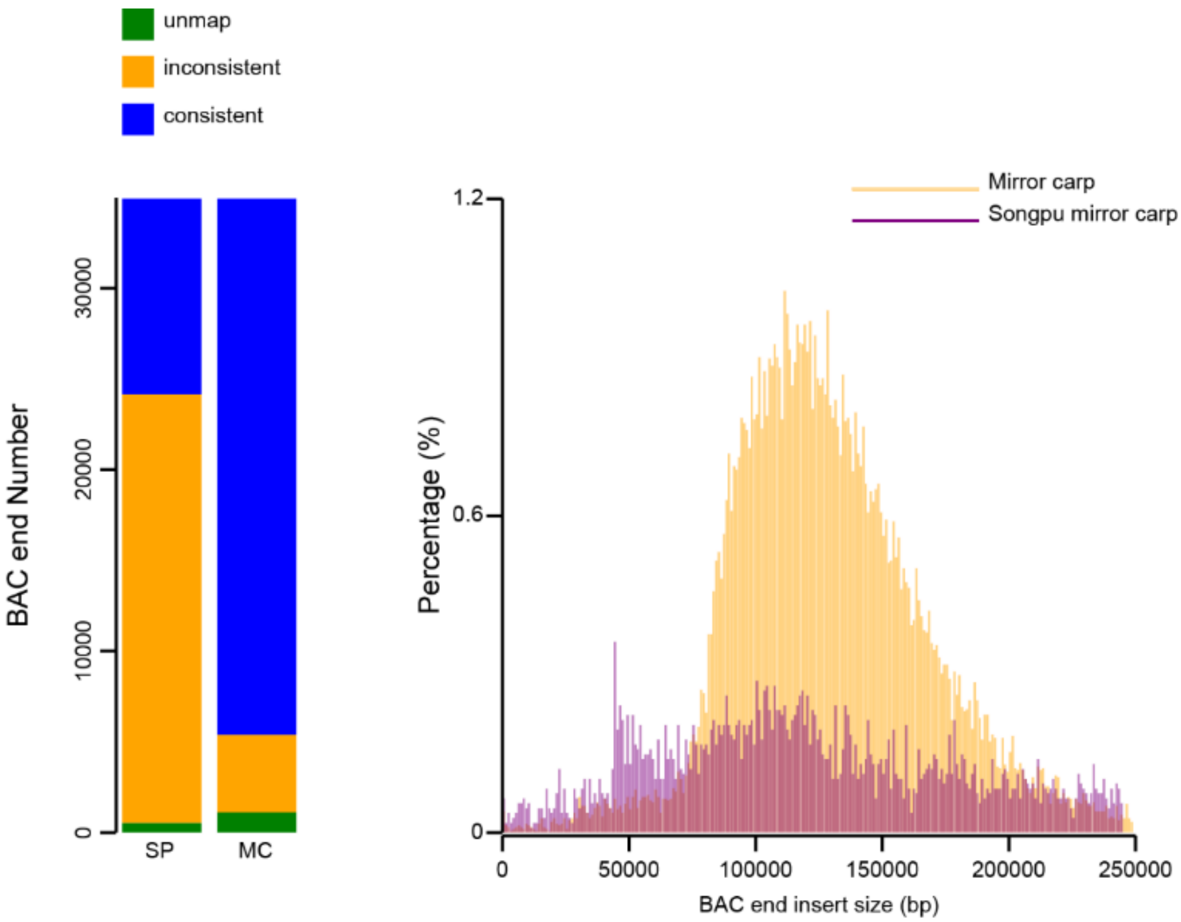

**Supplementary Figure 4. Circos plot of three common carp genomes.**

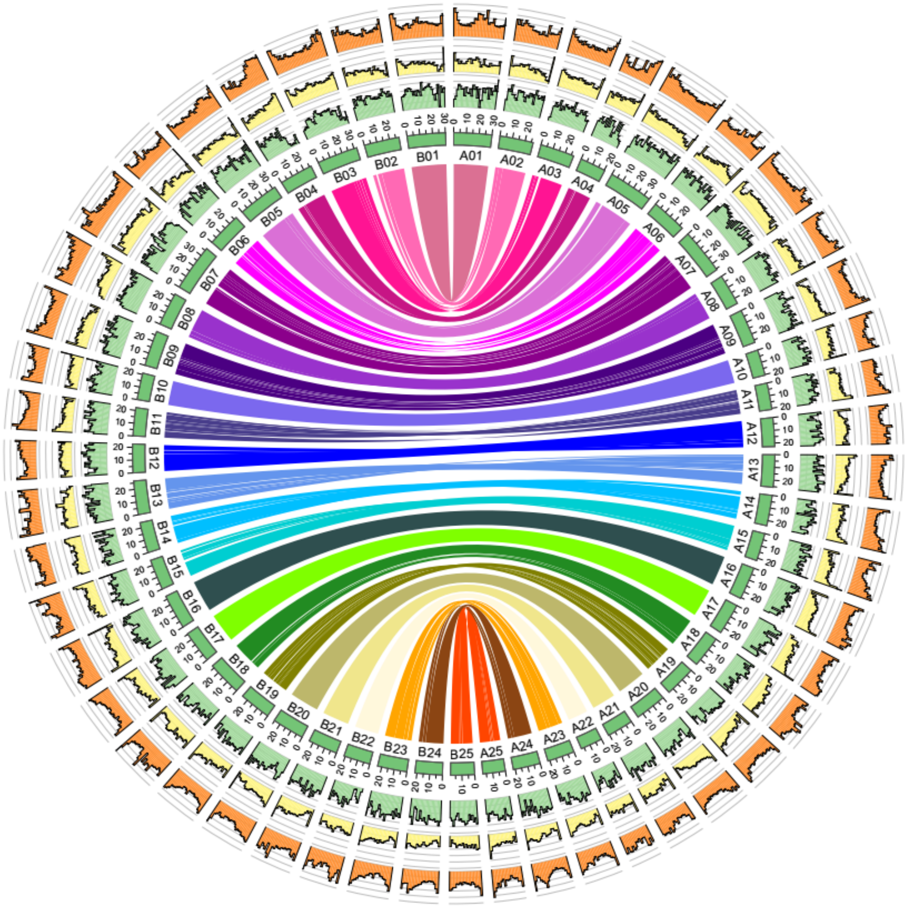

**(A) Yellow River Carp**

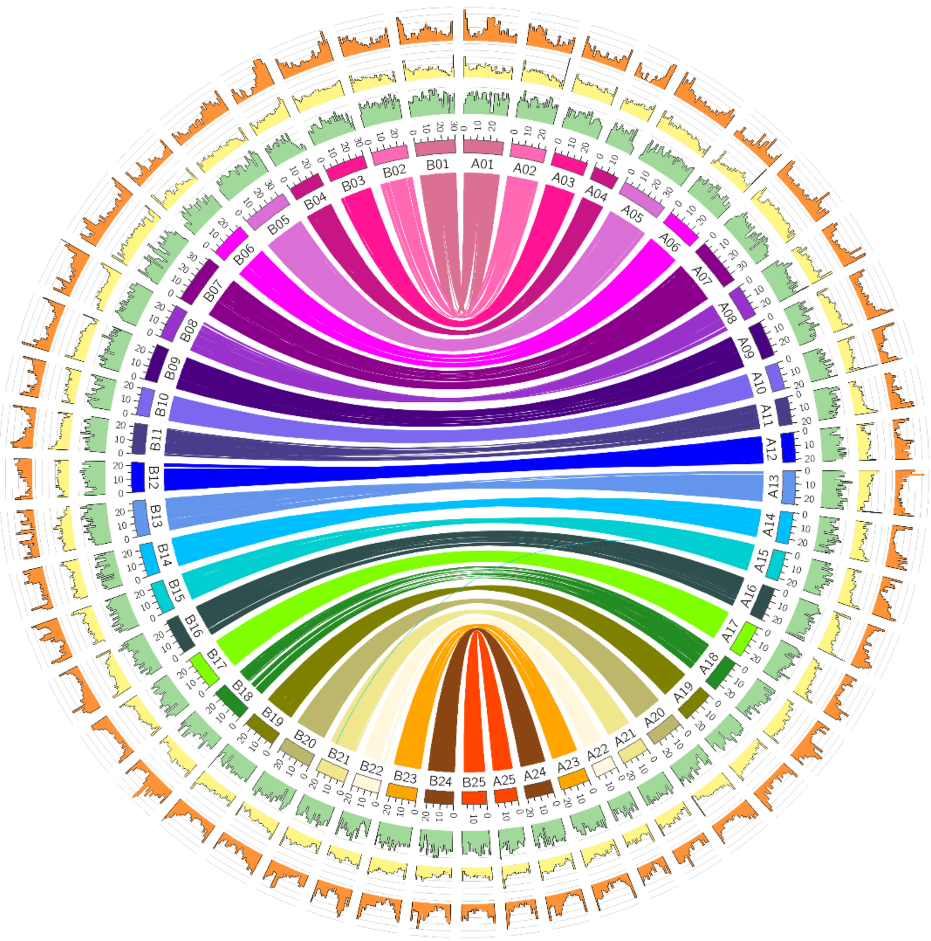

**(B) Hebao Red Carp**

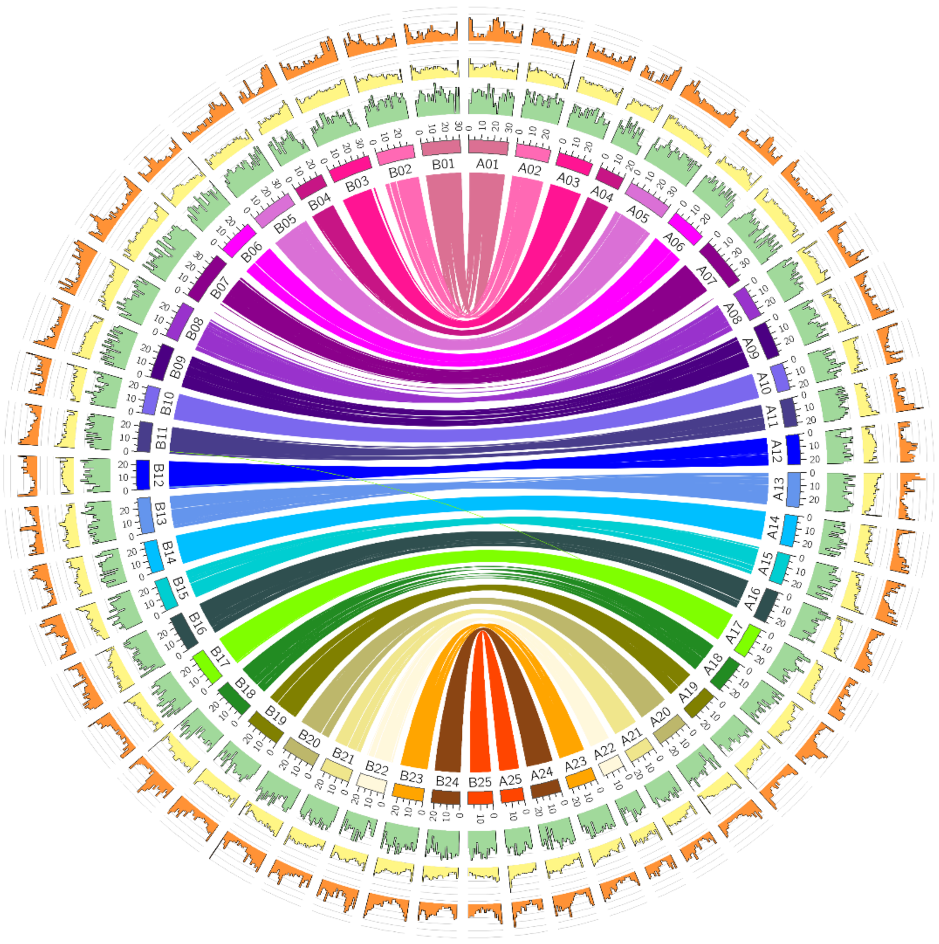

**(C) German Mirror Carp**

Note: Circos was used to plot repeat content within a 10-kb sliding window (the outermost plot, orange), GC content within a 10-kb sliding window (yellow), gene distribution on each chromosome (middle plot, green), assembled chromosomes and maps of the common carp A genome and B genome (innermost plot).

**Supplementary Figure 5. Phylogenetic tree of closely related Cyprininae species based on RAG2 and COI genes.**

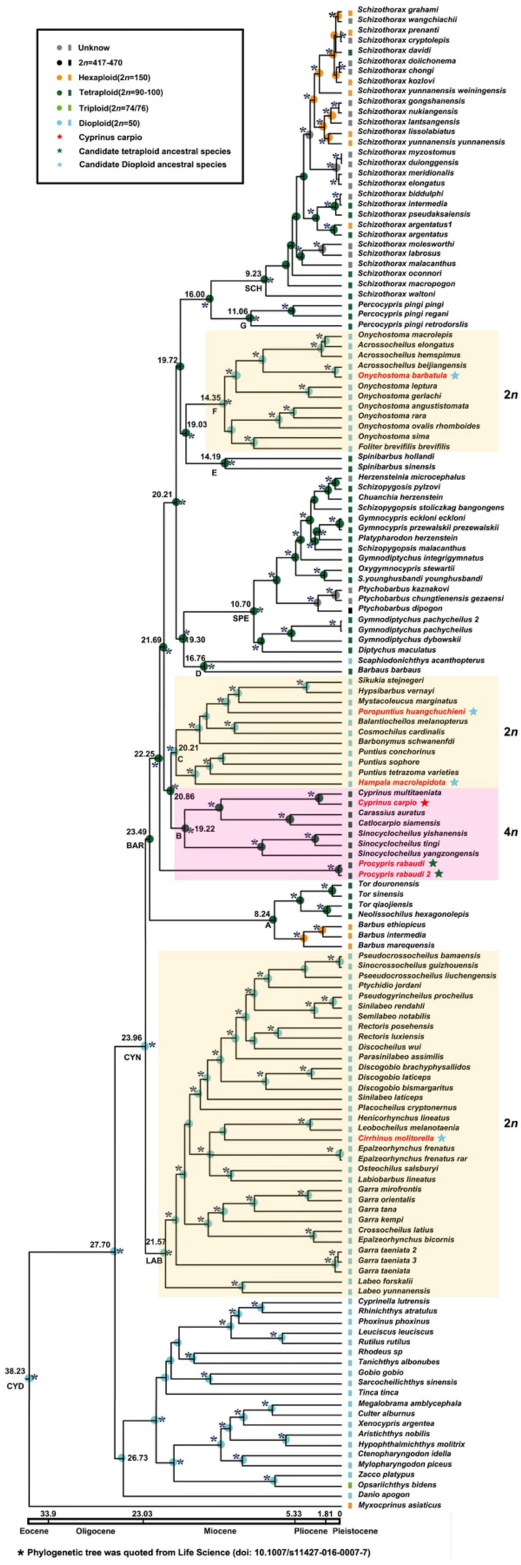

Figure was modified based on Figure 2 from Wang, et al, 2016 (DOI: 10.1007/s11427-016-0007-7).

**Supplementary Figure 6. Sampling sites of the sequenced candidate diploid progenitors.**

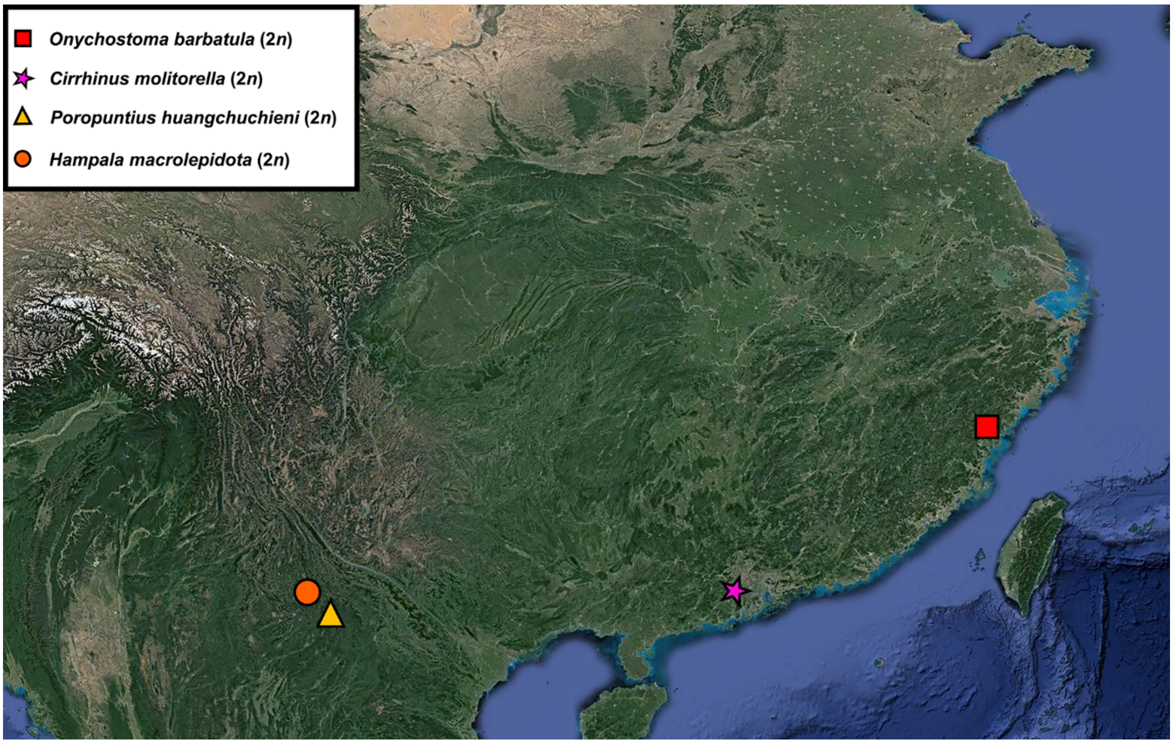

**Supplementary Figure 7. Phylogenetic tree based on 2071 conserved homoeologous gene pairs.**

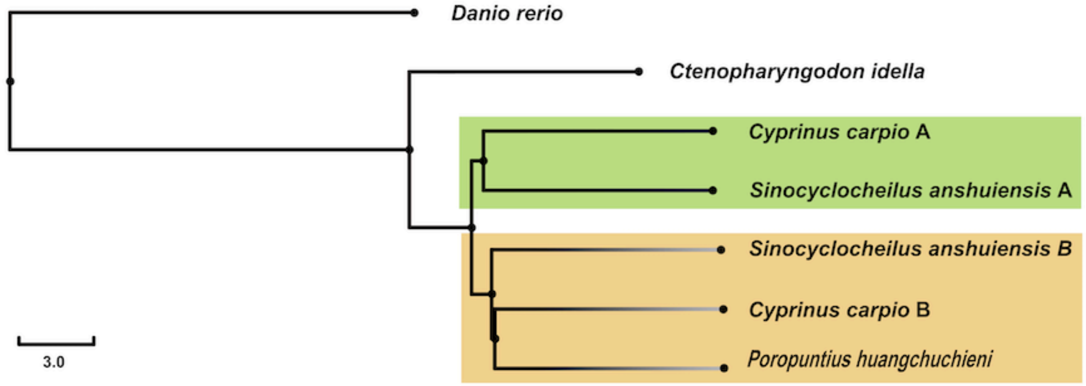

The 2071 phylogenetic trees were integrated using Bin-MPEST software.

**Supplementary Figure 8. Sequence of events for diversification (A), climate (B) and geology (C) in the region of the Qinghai-  Tibetan Plateau (QTP).**

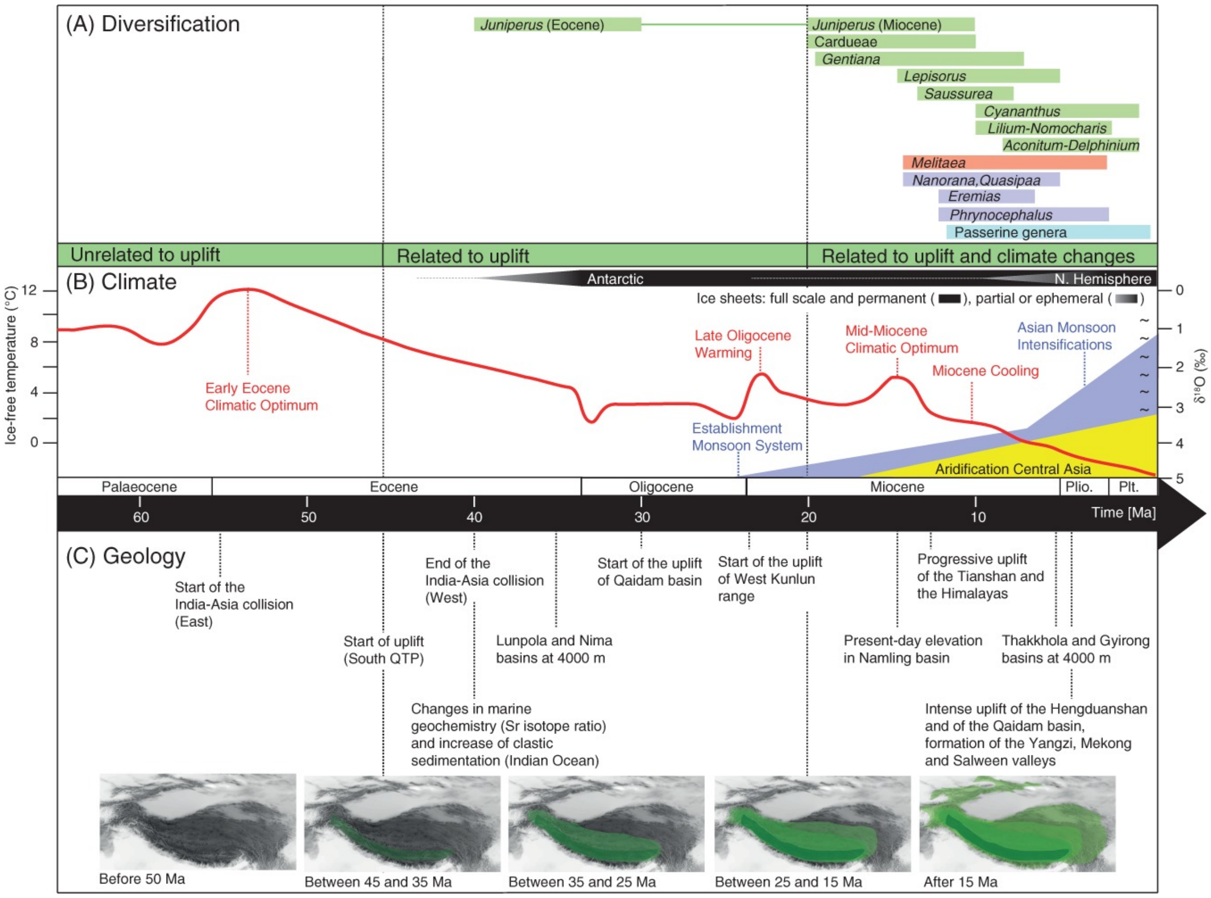

Note: The figure been previously published on Biological Reviews (Favre, Adrien , et al. 2015)

**Supplementary Figure 9. TE component in two subgenomes of common carp.**

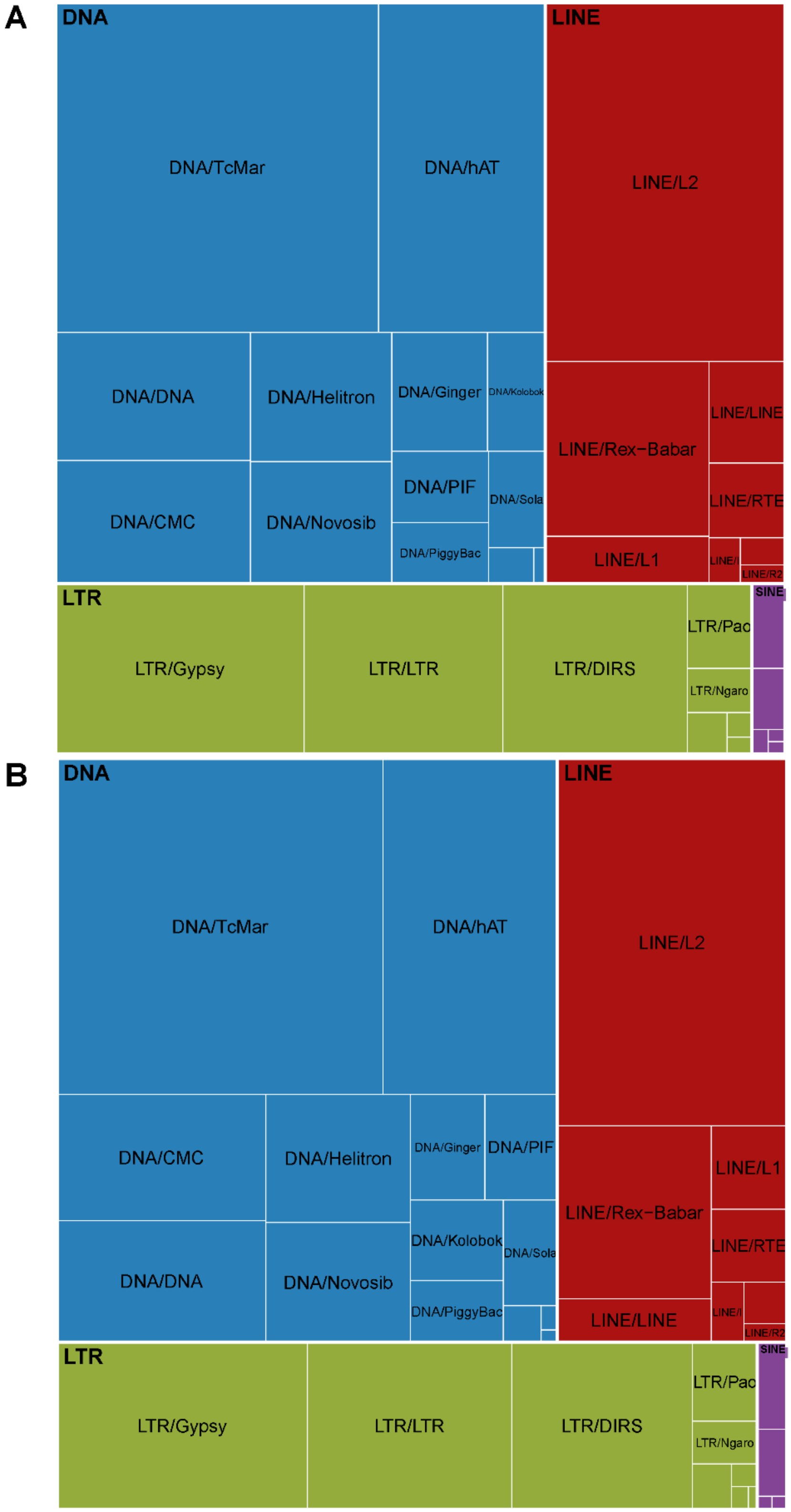

Note: **A**: subgenome A of common carp, **B**: subgenome B of common carp.

**Supplementary Figure 12. Proportions of duplicated and single-copy genes in tetraploid genome of *C.carpio*.**

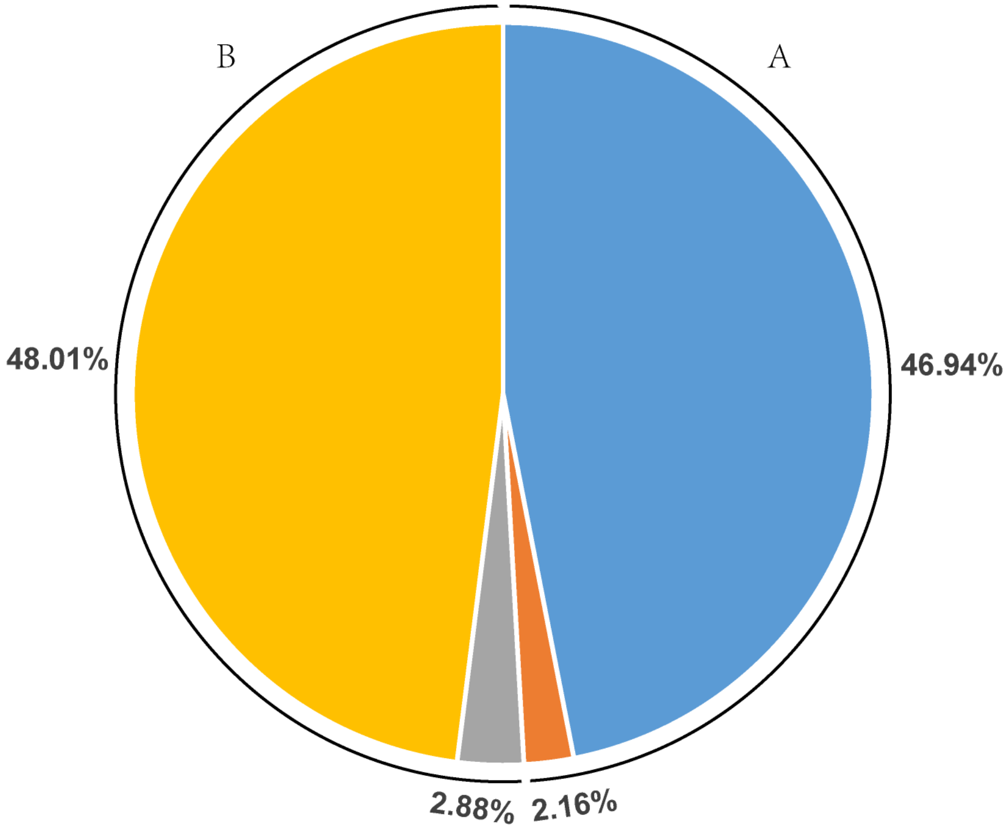

Note: A_paired: gene pairs in subgenome A, B_paired: gene pairs in subgenome B, A_singe: single genes in subgenome A, B_singe: single genes in subgenome B.

**Supplementary Figure 13. Typical structure variations between homoeologous chromosomes of two subgenomes**

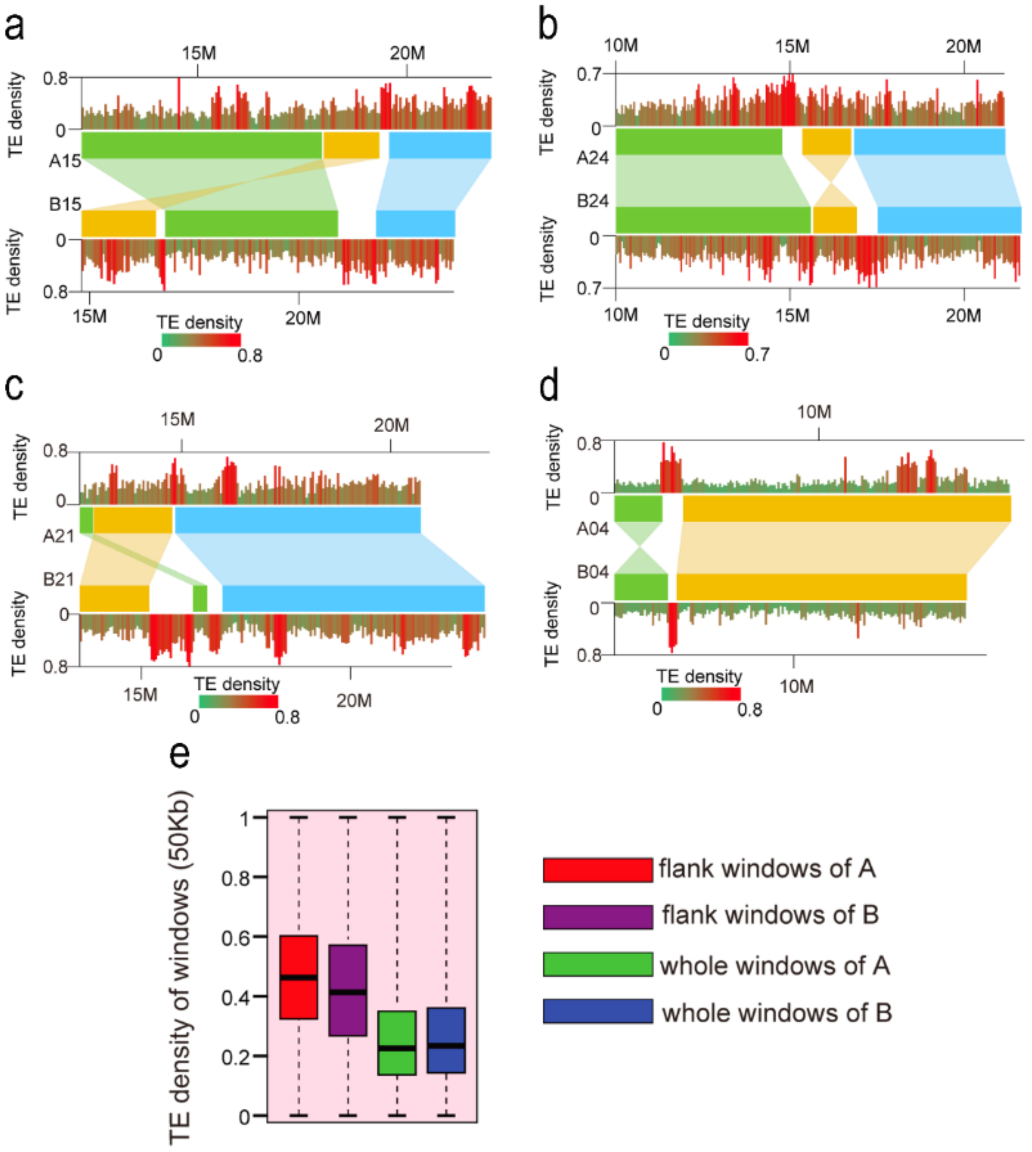

Note: Segmental translocation in the homoeologous chromosome A15/B15, A21/B21 (a,c) and segmental inversion in the homoeologous chromosome A24/B24, A04/B04 (b,d) of *C. carpio*; e, TE distribution in the flanking regions of the structure variations and chromosome-level TE distribution.

**Supplementary Figure 14. Expression patterns of total genes in the 12 tissues in *C. carpio*.**

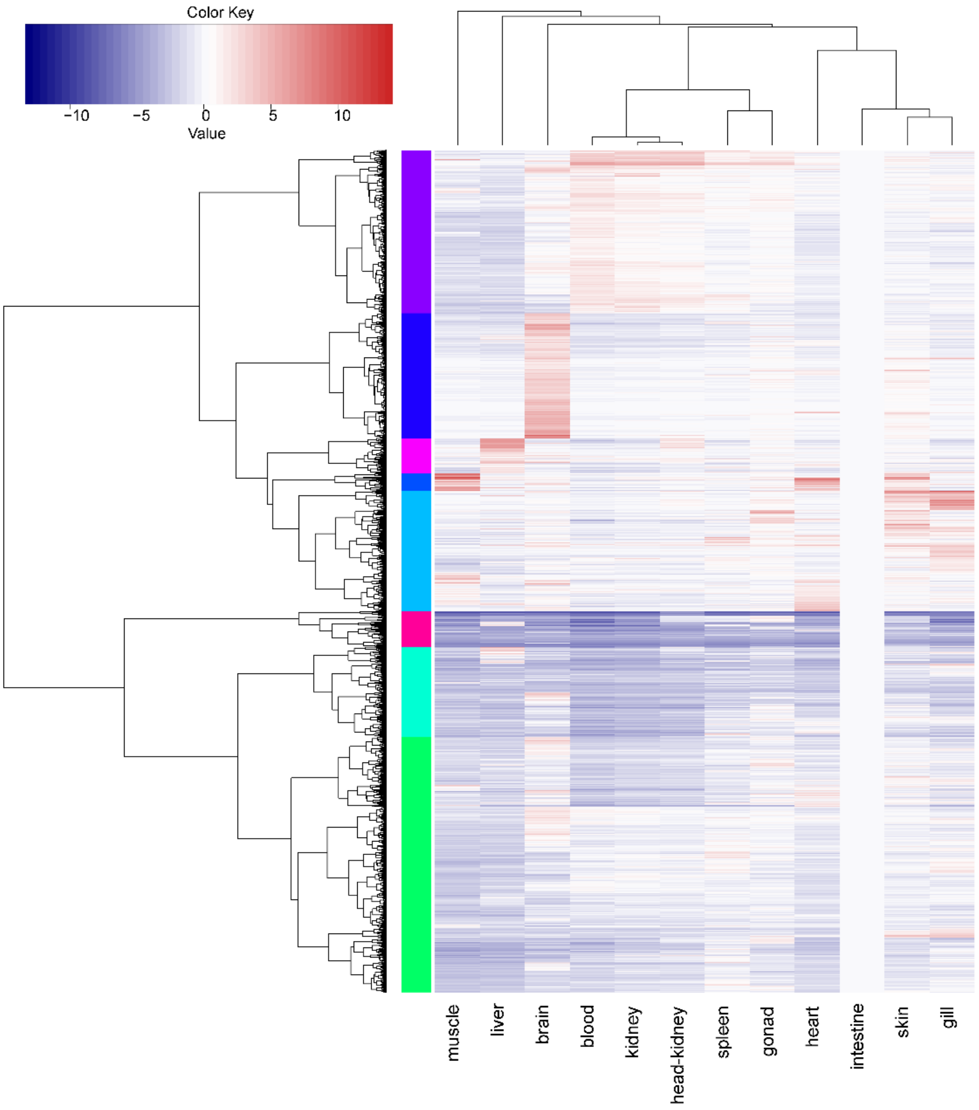

**Supplementary Figure 15. Clusters for 8291 homeologous gene pairs in the 12 tissues in *C. idella* and *C. carpio*.**

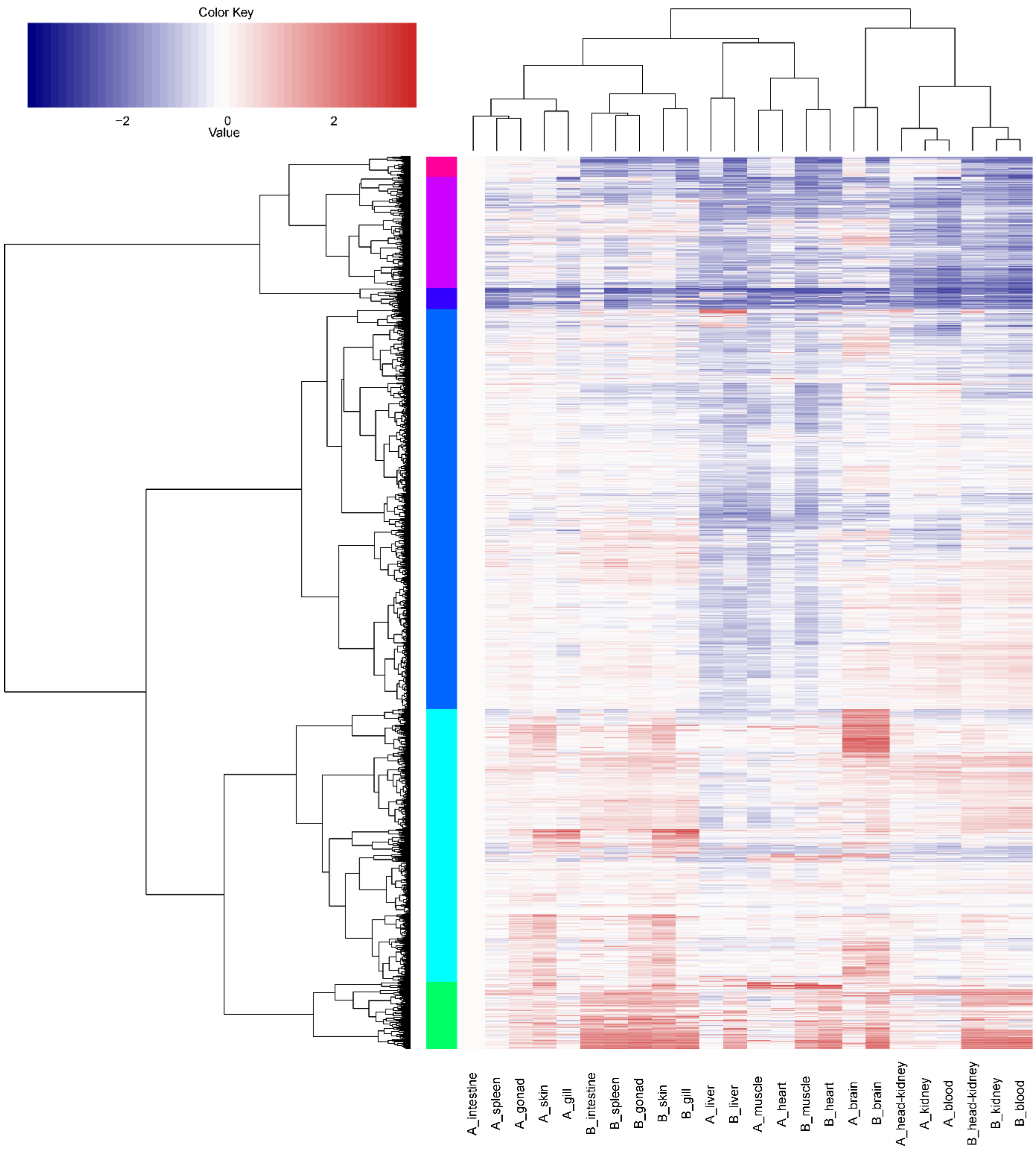

**Supplementary Figure 16. Expression patterns o.f *pdl1* and *acsl6* and genes in 12 tissues in common carp.**

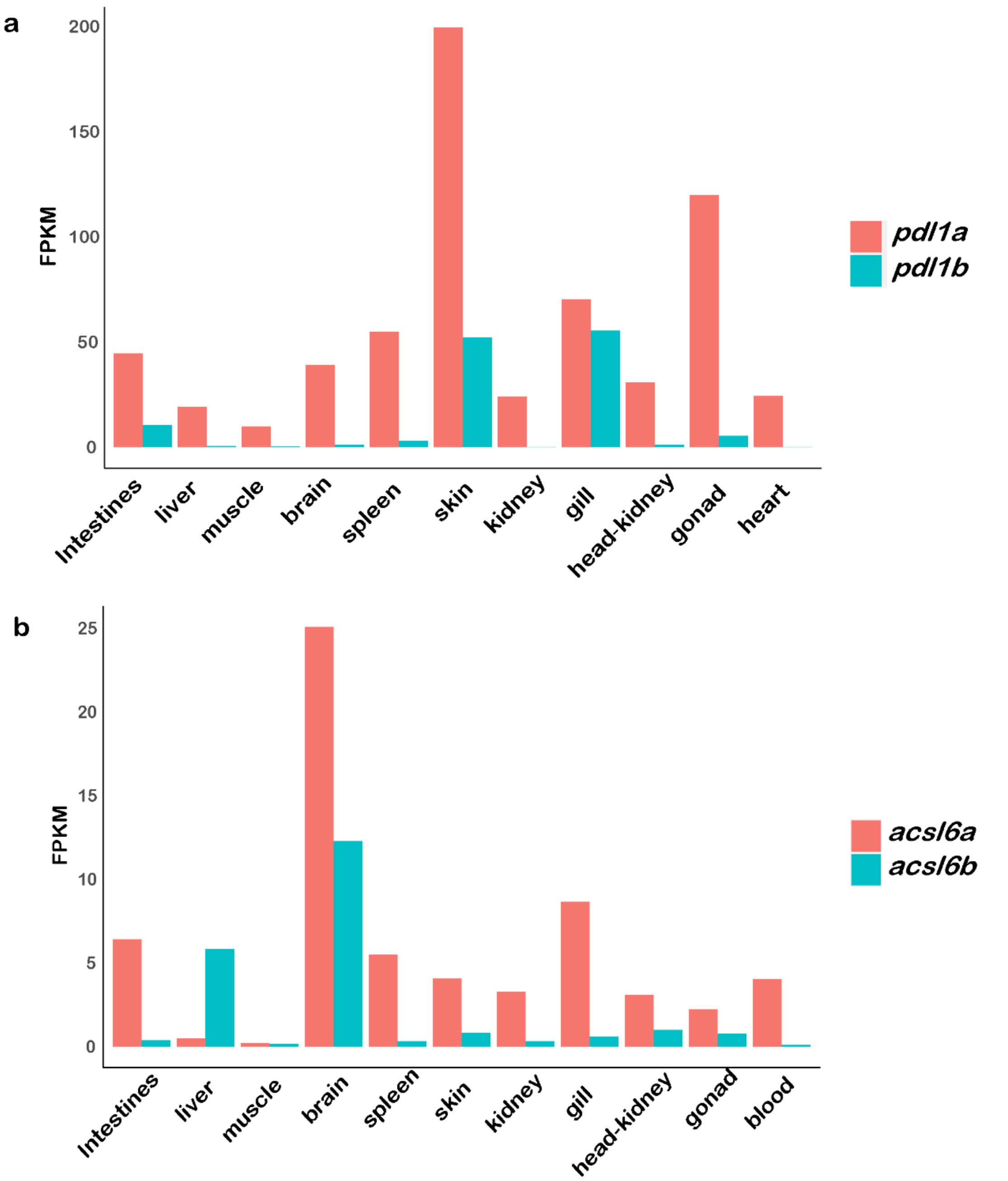

Note: **a**. Expression patterns of *pdl1* genes in 12 tissues; **b**. Expression patterns of *acsl6* genes in 12 tissues.

**Supplementary Figure 17. Asymmetric homoeologous expression patterns under biotic or abiotic stresses.**

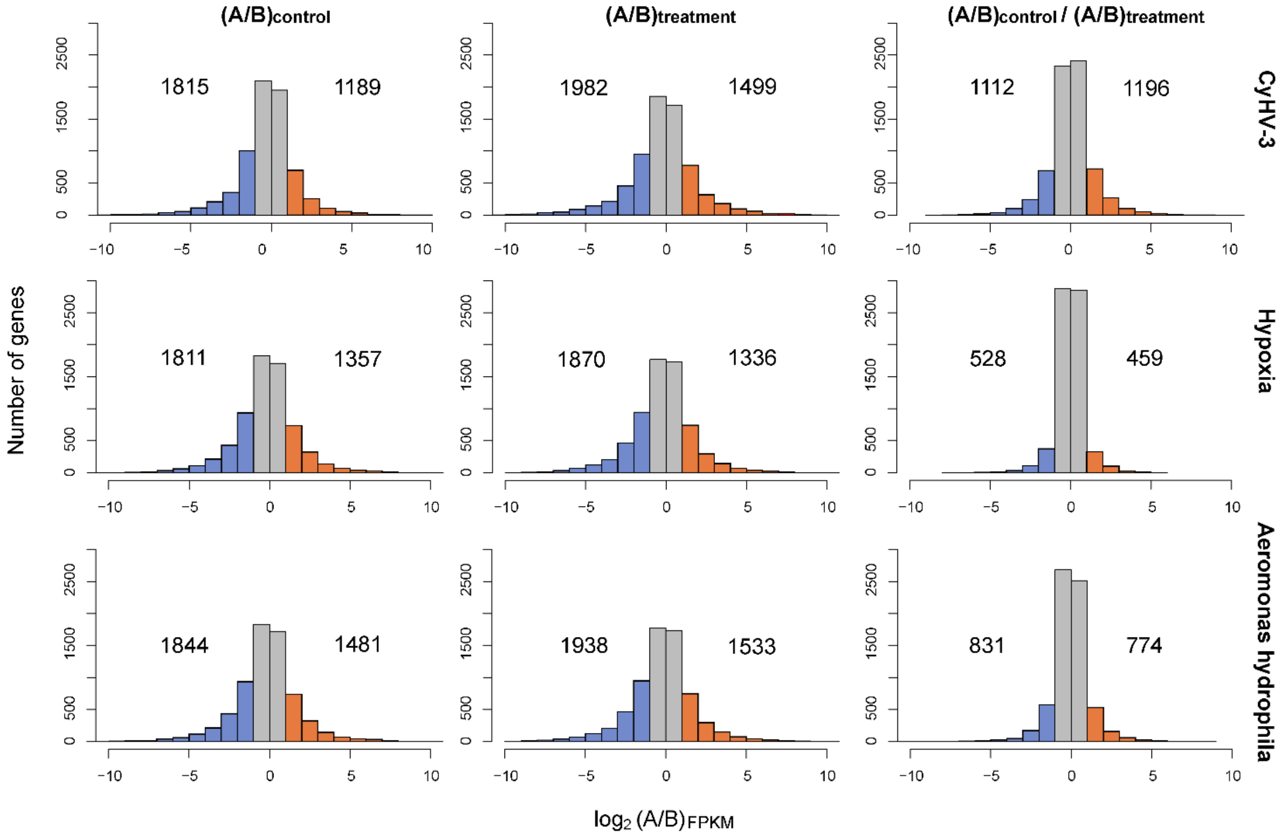

**Supplementary Figure 18. Methylation levels of asymmetrically expressed homoeologous genes in 12 tissues.**

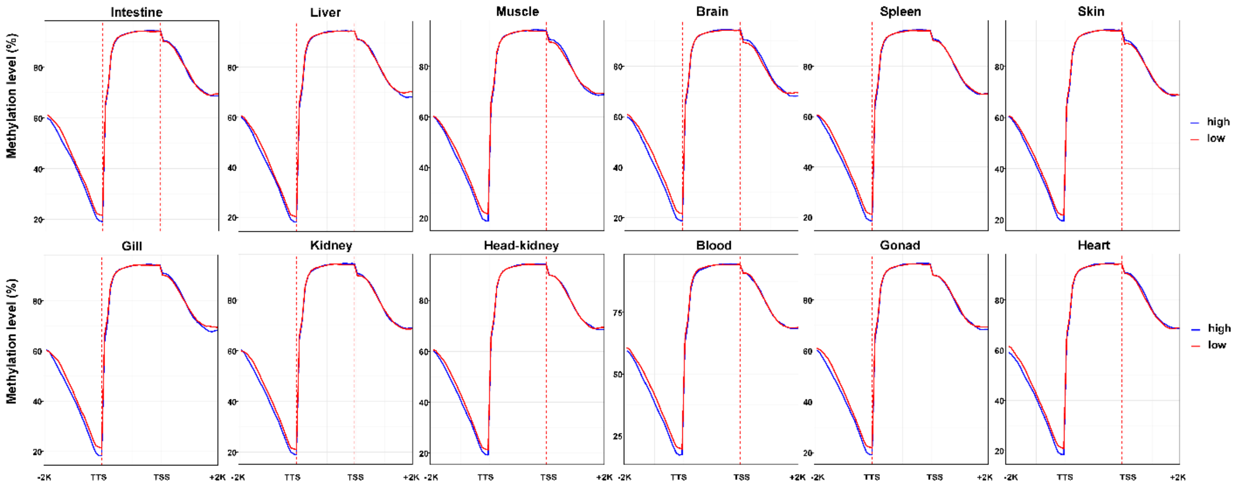

Note: High meas highly expressed genes, low meas lowly expressed genes.

**Supplementary Figure 19. CG methylation levels of divergent expressed homoeologous genes.**

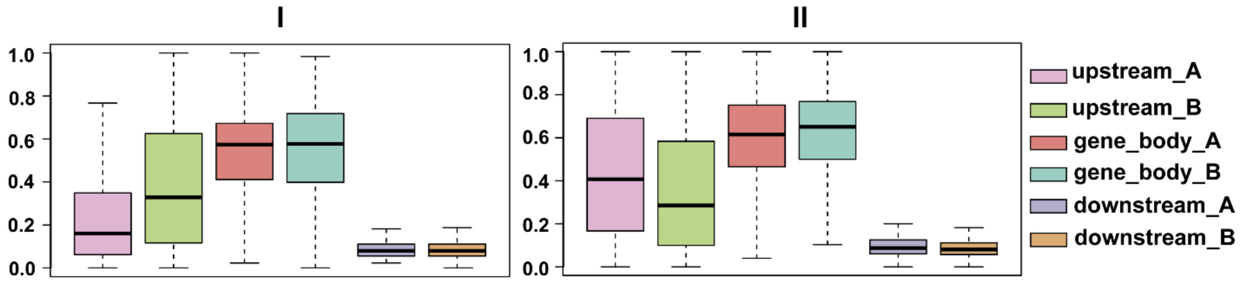

Note: I means homoeologous genes dominantly expressed in subgenome A but scilenced in subgenome B; II means homoeologous genes dominantly expressed in subgenome B but scilenced in subgenome A.

**Supplementary Figure 20. Genetic initrogression line (HB to YR) demonstrates skin color segregation.**

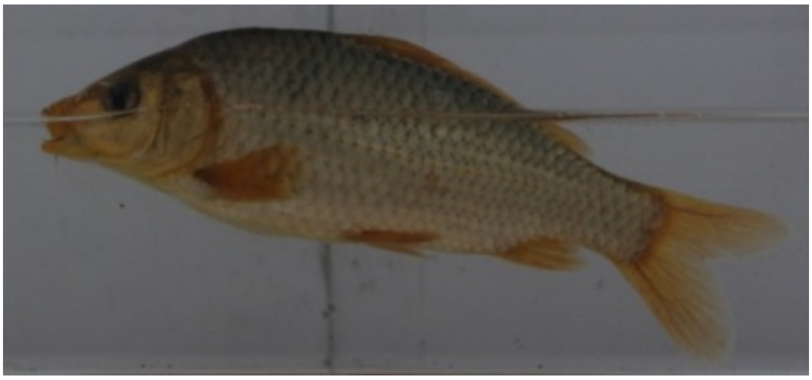

**(A) Reddish skin color**

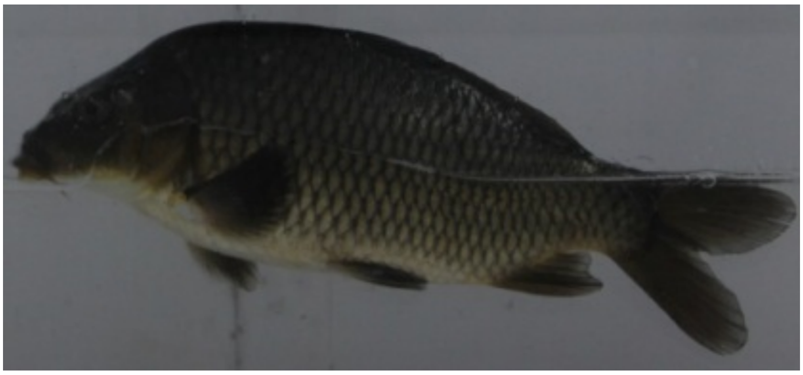

**(B) Wild-type skin color**

**Supplementary Figure 21. Genome wide association study in an introgression line with skin color segregation.**

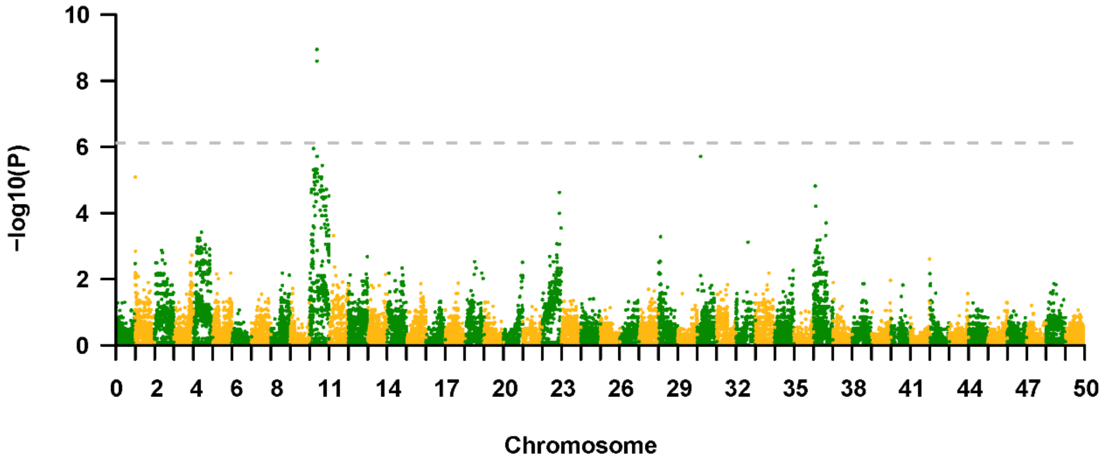

Note: Phenotype segregation of wild-type and red skin in the introgression line was 3:1. The gray dash line genome-wide significance threshold according to Bonferroni method (0.05/N= 6.12).

### Supplementary Tables

**Supplementary Table 1. Summaries of library construction and sequencing data of three common carp strains**

| Strain | Library Insert | Total base (Gbp) | Read length (bp) | Sequence coverage (X) |
|---|---|---|---|---|
| Hebao red carp | 230 bp | 77.69 | 150 | 48.56 |
| | 350 bp | 53.51 | 150 | 33.44 |
| | 230 bp | 47.47 | 150 | 29.67 |
| | 2 Kb | 27.96 | 150 | 17.48 |
| | 5 Kb | 32.56 | 125 | 20.35 |
| | 10 Kb | 46.22 | 125 | 28.89 |
| | 20 Kb | 13.24 | 125 | 8.28 |
| | Total | 298.65 | | 186.66 |
| Yellow River carp | 250 bp | 89.88 | 150 | 56.18 |
| | 500 bp | 77.01 | 150 | 48.13 |
| | 1.2 Kb | 33.28 | 150 | 20.80 |
| | 2.4 Kb | 33.70 | 150 | 21.06 |
| | 5 Kb | 38.35 | 150 | 23.96 |
| | 10 Kb | 34.54 | 150 | 21.59 |
| | 15 Kb | 18.41 | 150 | 11.51 |
| | 20 Kb | 13.94 | 150 | 8.71 |
| | Total | 339.11 | | 211.94 |
| Mirror carp | 250 bp | 89.11 | 150 | 59.80 |
| | 500 bp | 73.54 | 150 | 49.36 |
| | 2 Kb | 58.85 | 150 | 39.50 |
| | 5 Kb | 37.18 | 150 | 24.95 |
| | 10 Kb | 35.69 | 150 | 23.96 |
| | 15 Kb | 17.25 | 150 | 11.58 |
| | 20 Kb | 18.77 | 150 | 12.59 |
| | Total | 330.39 | | 221.74 |

**Supplementary Table 2. The genome assembly statistics of three common carp strains**

| Strains | | Length (bp) | | Number | |
|---|---|---|---|---|---|
| | | Contig | Scaffold | Contig | Scaffold |
| Hebao red carp | Total | 1,409,910,433 | 1,460,461,589 | 355,804 | 262,449 |
| | Max | 207,110 | 6,571,350 | - | - |
| | >=2000 bp | - | - | 87,986 | 9,135 |
| | N50 | 20,684 | 923,370 | 19,142 | 393 |
| | N60 | 15,917 | 629,080 | 26,914 | 589 |
| | N70 | 11,609 | 420,074 | 37,264 | 873 |
| | N80 | 7,300 | 205,075 | 52,431 | 1,362 |
| | N90 | 2,325 | 22,238 | 84,081 | 3,290 |
| Yellow River carp | Total | 1,392,182,197 | 1,424,610,504 | 254,656 | 144,810 |
| | Max | 217,672 | 10,558,934 | - | - |
| | >=2000 bp | - | - | 89,155 | 8,761 |
| | N50 | 21,811 | 1,706,244 | 18,518 | 211 |
| | N60 | 17,102 | 1,268,502 | 25,724 | 308 |
| | N70 | 12,799 | 878,940 | 35,132 | 444 |
| | N80 | 8,734 | 424,730 | 48,213 | 674 |
| | N90 | 4,302 | 97,382 | 70,189 | 1,331 |
| Mirror carp | Total | 1,359,861,568 | 1,415,500,317 | 76,948 | 18,344 |
| | Max | 409,856 | 14,180,042 | - | - |
| | >=2000 bp | - | - | 47,212 | 5,069 |
| | N50 | 52,141 | 3,465,820 | 7,457 | 107 |
| | N60 | 40,549 | 2,472,690 | 10,411 | 157 |
| | N70 | 30,524 | 1,769,284 | 14,280 | 224 |
| | N80 | 20,903 | 1,135,885 | 19,641 | 324 |
| | N90 | 11,395 | 421,233 | 28,250 | 520 |

**Supplementary Table 3. Summaries of genetic linkage maps of three common carp strains.**

| species | snp | locus | length (cM) | average marker interval (cM) | average locus interval (cM) |
|---|---|---|---|---|---|
| YR | 28194 | 14146 | 10596 | 0.38 | 0.75 |
| HB | 29019 | 16183 | 21964 | 0.76 | 1.36 |
| GM | 32160 | 23936 | 19425 | 0.60 | 0.81 |

**Supplementary Table 4. Statistics of three integrated genome assemblies.**

| Species | Genome size (bp) | Anchored genome size (bp) | Contige N50 (bp) | Scaffold N50 (bp) | Gene content | Anchored gene content | Anchored genome ratio | Anchored gene ratio |
|---|---|---|---|---|---|---|---|---|
| YR | 1,424,950,904 | 1,261,167,219 | 21,811 | 1,706,244 | 44626 | 42,303 | 88.51% | 94.79% |
| HB | 1,510,402,370 | 1,241,545,686 | 20,684 | 923,370 | 44269 | 41,569 | 82.20% | 93.90% |
| GM | 1,416,103,994 | 1,304,360,463 | 52,141 | 3,465,820 | 44758 | 43,577 | 92.11% | 97.36% |

**Supplementary Table 5. Completeness assessment of three common carp genomes.**

| Species | complete | | complete + partial | |
|---|---|---|---|---|
| | proteins | %completeness | proteins | %completeness |
| HB | 216 | 87.1 | 243 | 97.98 |
| YR | 216 | 87.1 | 245 | 98.79 |
| GM | 215 | 86.69 | 245 | 98.79 |

**Supplementary Table 6. Gene region coverage assessed based on ESTs in three common carp genome assemblies**

| Strain | Dataset | Number | Total length (bp) | Sequences Covered by assembly (%) | With >90% sequences In one scaffold | | With >50% sequences In one scaffold | |
|---|---|---|---|---|---|---|---|---|
| | | | | | Number | Percent (%) | Number | Percent (%) |
| HB | >0 bp | 13,198 | 7,679,659 | 99.86 | 12,433 | 94.20 | 13,135 | 99.52 |
| | >200 bp | 13,198 | 7,679,659 | 99.86 | 12,433 | 94.20 | 13,135 | 99.52 |
| | >500 bp | 4,279 | 5,036,033 | 99.91 | 3,942 | 92.12 | 4,250 | 99.32 |
| | >1 Kb | 1,793 | 3,310,438 | 99.89 | 1,621 | 90.41 | 1,778 | 99.16 |
| | >2 Kb | 517 | 1,528,476 | 99.81 | 458 | 88.59 | 511 | 98.84 |
| YR | >0 bp | 23,352 | 12,465,030 | 99.26 | 21,647 | 92.70 | 22,996 | 98.48 |
| | >200 bp | 23,352 | 12,465,030 | 99.26 | 21,647 | 92,70 | 22,996 | 98.48 |
| | >500 bp | 7,285 | 7,714,476 | 99.64 | 6,829 | 93.74 | 7,225 | 99.18 |
| | >1 Kb | 2,764 | 4,589,503 | 99.89 | 2,601 | 94.10 | 2,749 | 99.46 |
| | >2 Kb | 583 | 1,590,318 | 99.83 | 547 | 93.83 | 577 | 98.97 |
| GM | >0 bp | 23,409 | 13,865,382 | 99.20 | 21,770 | 93.00 | 22,998 | 98.24 |
| | >200 bp | 23,409 | 13,865,382 | 99.20 | 21,770 | 93.00 | 22,998 | 98.24 |
| | >500 bp | 8,077 | 9,321,683 | 99.68 | 7,539 | 93.34 | 7,992 | 98.95 |
| | >1 Kb | 3,484 | 6,119,622 | 99.94 | 3,287 | 94.35 | 3,464 | 99.43 |
| | >2 Kb | 894 | 2,522,202 | 99.89 | 835 | 93.40 | 887 | 99.22 |

**Supplementary Table 7. Transposable elements in common carp genomes.**

| Repeat class | Hebao red carp | | Yellow River carp | | German mirror carp | |
|---|---|---|---|---|---|---|
| | length | rate (%) | length | rate (%) | length | rate (%) |
| DNA/hAT | 46,187,953 | 3.06 | 42,708,696 | 2.99 | 39,955,117 | 2.82 |
| DNA/TcMar-Tc1 | 76,687,355 | 5.08 | 71,927,799 | 5.05 | 74,796,224 | 5.28 |
| DNA/other | 86,767,046 | 5.74 | 84,504,475 | 5.93 | 78,029,897 | 5.51 |
| LINE/L2 | 64,392,604 | 4.26 | 63,835,670 | 4.48 | 60,937,527 | 4.31 |
| LINE/other | 45,027,580 | 2.98 | 41,679,366 | 2.92 | 39,509,774 | 2.79 |
| LTR/Gypsy | 40,777,238 | 2.69 | 36,324,585 | 2.55 | 36,049,641 | 2.55 |
| LTR/other | 66,297,715 | 4.39 | 59,795,318 | 4.19 | 58,439,616 | 4.13 |
| RC/Helitron | 14,231,312 | 0.94 | 13,432,925 | 0.94 | 13,134,833 | 0.93 |
| Unknown | 66,929,472 | 4.43 | 54,729,884 | 3.84 | 47,539,485 | 3.36 |
| Other | 50,576,220 | 3.34 | 50,043,491 | 3.51 | 45,707,911 | 3.22 |
| Total | 557,874,495 | 36.94 | 518,982,209 | 36.42 | 494,100,025 | 34.89 |

**Supplementary Table 8. Gene prediction in common carp genome.**

| Species | Genome length (bp) | Total gene length (bp) | Gene content | Exon content | Exon number per gene | Gene density | Average gene length (bp) |
|---|---|---|---|---|---|---|---|
| YR | 1,424,950,904 | 616,471,225 | 44,626 | 426,870 | 9.57 | 0.43 | 13,814.17 |
| HB | 1,510,402,370 | 602,203,106 | 44,269 | 418,078 | 9.44 | 0.40 | 13,603.27 |
| GM | 1,416,103,994 | 626,902,020 | 44,758 | 428,776 | 9.58 | 0.44 | 14,006.48 |

**Supplementary Table 9. Gene annotation in common carp genome.**

| Database | Hebao red carp | Yellow River carp | German mirror carp |
|---|---|---|---|
| Swiss-prot | 41,019 | 40,710 | 40,790 |
| | (92.6%) | (91.2%) | (91.1%) |
| NR | 42,893 | 42,909 | 43,016 |
| | (96.9%) | (96.2%) | (96.1%) |
| KEGG | 37,258 | 37,108 | 37,210 |
| | (84.1%) | (83.2%) | (83.1%) |
| GO | 31,004 | 31,041 | 31,154 |
| | (70%) | (69.6%) | (69.6%) |
| Pfam | 37,713 | 37,797 | 37,859 |
| | (85.2%) | (84.7%) | (84.6%) |
| InterPro | 40,872 | 40,721 | 40,772 |
| | (92.3%) | (91.2%) | (91.1%) |
| Annotated | 42,916 | 42,916 | 43,055 |
| | (96.9%) | (96.2%) | (96.2%) |

**Supplementary Table 10. Accession number of *rag2* gene used in this study.**

| Species | Gene | Accession Number |
|---|---|---|
| *Acrossocheilus hemispinus* | *rag2* | DQ366986 |
| *Onychostoma barbatula* | *rag2* | DQ366964 |
| *Onychostoma angustistomata* | *rag2* | HQ235896 |
| *Hampala macrolepidota* | *rag2* | DQ366965 |
| *Puntius tetrazoma* | *rag2* | DQ366938 |
| *Puntius sophore* | *rag2* | HQ235899 |
| *Mystacoleucus marginatus)* | *rag2* | HQ235894 |
| *Poropuntius huangchuchieni* | *rag2* | DQ366952 |
| *Sinocyclocheilus anshuiensis* | *rag2a* | XM_016249750.1 |
| *Sinocyclocheilus anshuiensis* | *rag2b* | XM_016466752.1 |
| *Garra tana* | *rag2* | HQ235879 |
| *Cirrhinus molitorella* | *rag2* | DQ366959 |
| *Discogobio laticeps* | *rag2* | DQ366949 |
| *Sinilabeo laticeps* | *rag2* | HQ235885 |
| *Ctenopharyngodon idella* | *rag2* | DQ366996 |
| *Danio apogon* | *rag2* | U71094 |

**Supplementary Table 11. Genome sequencing and assembly of diploid species.**

| Species | | Length (bp) | | Number | |
|---|---|---|---|---|---|
| | | Contig** | Scaffold | Contig** | Scaffold |
| *Poropuntius huangchuchieni* | Total | 846,673,898 | 868,595,308 | 1,667,071 | 1,431,194 |
| | Max | 76,402 | 140,771 | - | - |
| | Number>=2000 | - | - | 82,623 | 84,023 |
| | N50 | 1,574 | 2,176 | 109,206 | 76,628 |
| *Cirrhinus molititorella* | Total | 953,783,459 | 985,712,601 | 1,271,496 | 976,377 |
| | Max | 93,971 | 145,483 | - | - |
| | Number>=2000 | - | - | 139,665 | 139,179 |
| | N50 | 2,444 | 3,824 | 109,206 | 76,628 |

| Species | Raw Reads | Raw bases | Clean Reads | Clean base |
|---|---|---|---|---|
| *Onychostoma barbatula* | 214,371,286 | 64,311,385,800 | 183,276,646 | 54,982,993,800 |
| *Hampala macrolepidota* | 479,970,766 | 71,995,614,900 | 33,285,511 | 4,992,826,688 |
| *Procyris rabaudi* | 37,652,816 | 5,647,922,438 | 35,398,784 | 5,309,817,525 |

**Supplementary Table 12. Chromosome nomenclature of *Cyprinus carpio* according to *Danio rerio*.** Chromosomes in *Cyprinus carpio* were named as A and B based on the progenitor genome.

| *Danio rerio* | *Cyprinus carpio* | |
|---|---|---|
| | subgenome A | subgenome B |
| LG1 | A01 | B01 |
| LG2 | A02 | B02 |
| LG3 | A03 | B03 |
| LG4 | A04 | B04 |
| LG5 | A05 | B05 |
| LG6 | A06 | B06 |
| LG7 | A07 | B07 |
| LG8 | A08 | B08 |
| LG9 | A09 | B09 |
| LG10 | A10 | B10 |
| LG11 | A11 | B11 |
| LG12 | A12 | B12 |
| LG13 | A13 | B13 |
| LG14 | A14 | B14 |
| LG15 | A15 | B15 |
| LG16 | A16 | B16 |
| LG17 | A17 | B17 |
| LG18 | A18 | B18 |
| LG19 | A19 | B19 |
| LG20 | A20 | B20 |
| LG21 | A21 | B21 |
| LG22 | A22 | B22 |
| LG23 | A23 | B23 |
| LG24 | A24 | B24 |
| LG25 | A25 | B25 |

**Supplementary Table 13. Gene content of each chromosome of C. carpio.**

| Chr | Gene content | | | Gene_density | | | Average_gene_length | | |
|---|---|---|---|---|---|---|---|---|---|
| | YR | HB | GM | YR | HB | GM | YR | HB | GM |
| A01 | 945 | 947 | 953 | 0.44 | 0.44 | 0.43 | 13849.7 | 13799.4 | 14096.6 |
| A02 | 980 | 931 | 1002 | 0.51 | 0.49 | 0.51 | 13825.4 | 13834.2 | 13754 |
| A03 | 1091 | 1038 | 1086 | 0.45 | 0.45 | 0.45 | 11035.7 | 11136.6 | 11445.2 |
| A04 | 613 | 588 | 611 | 0.45 | 0.46 | 0.45 | 15092.3 | 14709.9 | 14978.7 |
| A05 | 1213 | 1177 | 1231 | 0.51 | 0.48 | 0.51 | 14622.4 | 14286.9 | 14662.7 |
| A06 | 950 | 927 | 947 | 0.49 | 0.48 | 0.48 | 14405.1 | 14282.7 | 14002.6 |
| A07 | 1079 | 1089 | 1138 | 0.47 | 0.46 | 0.44 | 15577.4 | 14652.5 | 14856.2 |
| A08 | 923 | 895 | 937 | 0.52 | 0.54 | 0.52 | 13863.7 | 14554.4 | 14181.6 |
| A09 | 764 | 752 | 778 | 0.47 | 0.47 | 0.48 | 16574.2 | 16853.7 | 17192 |
| A10 | 767 | 741 | 776 | 0.49 | 0.50 | 0.48 | 13079.6 | 13572 | 13457.3 |
| A11 | 697 | 686 | 733 | 0.50 | 0.49 | 0.50 | 14917.2 | 14862.8 | 15335.7 |
| A12 | 769 | 723 | 777 | 0.48 | 0.47 | 0.46 | 14219.3 | 14446.8 | 13954.4 |
| A13 | 847 | 796 | 862 | 0.51 | 0.51 | 0.48 | 15290.7 | 16336.2 | 15179.1 |
| A14 | 715 | 698 | 733 | 0.38 | 0.40 | 0.40 | 12972.6 | 13489.2 | 13469.4 |
| A15 | 739 | 732 | 759 | 0.49 | 0.48 | 0.46 | 14543.2 | 14456.6 | 14572.8 |
| A16 | 967 | 935 | 982 | 0.47 | 0.47 | 0.47 | 12521.3 | 12547.6 | 12290.9 |
| A17 | 806 | 811 | 821 | 0.50 | 0.48 | 0.48 | 15216.6 | 14672.9 | 14575.2 |
| A18 | 689 | 683 | 716 | 0.43 | 0.44 | 0.44 | 15358.7 | 15991.4 | 15952.6 |
| A19 | 861 | 837 | 869 | 0.45 | 0.45 | 0.45 | 12686.2 | 12991.6 | 12488 |
| A20 | 873 | 822 | 857 | 0.49 | 0.47 | 0.47 | 13896.1 | 13861.5 | 13805.2 |
| A21 | 784 | 774 | 800 | 0.51 | 0.51 | 0.50 | 13657.1 | 13476.3 | 12876.1 |
| A22 | 611 | 570 | 584 | 0.41 | 0.43 | 0.44 | 11726.8 | 12291.1 | 12997.9 |
| A23 | 819 | 805 | 834 | 0.49 | 0.49 | 0.48 | 12968.4 | 13060.7 | 12952.2 |
| A24 | 614 | 578 | 634 | 0.51 | 0.49 | 0.47 | 17828.3 | 17582.7 | 17042.1 |
| A25 | 656 | 637 | 658 | 0.48 | 0.48 | 0.49 | 13122.9 | 12919 | 13458.1 |
| Average_A | 830.88 | 806.88 | 843.12 | 0.48 | 0.47 | 0.47 | 14114.036 | 14186.748 | 14143.064 |

| | | | | | | | | | |
|---|---|---|---|---|---|---|---|---|---|
| B01 | 995 | 966 | 967 | 0.47 | 0.46 | 0.45 | 14263.9 | 14470.4 | 14123.8 |
| B02 | 1052 | 1022 | 1093 | 0.51 | 0.51 | 0.49 | 13480.6 | 13550.5 | 13257.7 |
| B03 | 1161 | 1099 | 1193 | 0.41 | 0.43 | 0.41 | 11366.1 | 11675.6 | 11368.1 |
| B04 | 686 | 649 | 698 | 0.37 | 0.41 | 0.43 | 14114.4 | 15165.3 | 14508.7 |
| B05 | 1188 | 1200 | 1204 | 0.52 | 0.52 | 0.51 | 14107.5 | 14594.5 | 13909.4 |
| B06 | 950 | 940 | 992 | 0.50 | 0.48 | 0.48 | 14214.2 | 13632.1 | 13684.5 |
| B07 | 1122 | 1114 | 1180 | 0.47 | 0.46 | 0.48 | 15579.5 | 15108.2 | 15561.6 |
| B08 | 949 | 920 | 1010 | 0.52 | 0.51 | 0.50 | 14472.3 | 14455.7 | 14727 |
| B09 | 790 | 773 | 793 | 0.48 | 0.47 | 0.49 | 17030.8 | 16696.7 | 16991.8 |
| B10 | 782 | 768 | 801 | 0.55 | 0.51 | 0.51 | 14770.6 | 13805.1 | 14124.9 |
| B11 | 739 | 736 | 759 | 0.51 | 0.50 | 0.49 | 15448.2 | 15651.1 | 15471.6 |
| B12 | 760 | 733 | 780 | 0.50 | 0.48 | 0.49 | 14850.3 | 14903.4 | 14809.5 |
| B13 | 827 | 836 | 878 | 0.53 | 0.51 | 0.50 | 17065.2 | 16614.4 | 16083.7 |
| B14 | 757 | 736 | 755 | 0.40 | 0.41 | 0.40 | 12742.7 | 13321.5 | 13406.6 |
| B15 | 784 | 742 | 786 | 0.48 | 0.49 | 0.47 | 14904.8 | 15469.6 | 14685.8 |
| B16 | 1003 | 973 | 1048 | 0.48 | 0.48 | 0.47 | 13192.4 | 13096 | 12661.7 |
| B17 | 824 | 810 | 831 | 0.50 | 0.49 | 0.50 | 15500 | 15393.5 | 15966 |
| B18 | 779 | 712 | 762 | 0.43 | 0.43 | 0.45 | 15079.9 | 15679.2 | 15797.3 |
| B19 | 879 | 863 | 914 | 0.46 | 0.45 | 0.41 | 13151.3 | 12950.7 | 12684.1 |
| B20 | 850 | 834 | 891 | 0.49 | 0.49 | 0.49 | 14693 | 14899 | 14832.1 |
| B21 | 842 | 802 | 846 | 0.49 | 0.48 | 0.49 | 13507.1 | 13717.4 | 13891.5 |
| B22 | 626 | 623 | 661 | 0.38 | 0.40 | 0.38 | 12667.2 | 12910.4 | 13193.4 |
| B23 | 860 | 836 | 900 | 0.51 | 0.50 | 0.49 | 13770.4 | 13469.8 | 13311.1 |
| B24 | 635 | 614 | 652 | 0.52 | 0.49 | 0.51 | 17652.8 | 17296.6 | 17471.7 |
| B25 | 691 | 676 | 705 | 0.52 | 0.50 | 0.49 | 14314.3 | 13712 | 13803.1 |
| Average_B | 861.24 | 839.08 | 883.96 | 0.48 | 0.47 | 0.47 | 14477.58 | 14489.548 | 14413.068 |
| Average_AB | 846.06 | 822.98 | 863.54 | 0.48 | 0.47 | 0.47 | 14295.808 | 14338.148 | 14278.066 |

**Supplementary Table 14. The statistics of gene structure of *C. carpio* and the comparison with other teleosts.**

| Gene Set | Genome assembly size (Mb) | No. genes | Mean CDS length | No. exons per gene | Mean exon size (bp) | Mean intron size (bp) |
|---|---|---|---|---|---|---|
| YR | 1,425 | 44,626 | 1,648 | 9.57 | 172 | 1,420 |
| HB | 1,510 | 44,269 | 1,641 | 9.44 | 174 | 1,417 |
| GM | 1,416 | 44,758 | 1,650 | 9.58 | 172 | 1,440 |
| *D. rerio* | 1,412 | 26,163 | 1,853 | 7.97 | 232 | 2,923 |
| *C. idellus* | 900 | 27,263 | 1,539 | 8.5 | 250 | 1,287 |
| *G.aculeatus* | 461 | 20,787 | 1,592 | 9.88 | 161 | 771 |
| *O.latipes* | 868 | 19,686 | 1,553 | 10.04 | 154 | 1,215 |
| *T.rubripes* | 393 | 18,523 | 1,617 | 10.69 | 151 | 600 |

Note: YR: Yellow River carp, HB: Hebao red carp, GM: German mirror carp.

**Supplementary Table 15. Gene present and loss categories based on distinguished A/B subgenome.** CID: *Ctenopharyngodon idella*; CCA: *C. carpio* A subgenome; CCB: *C. carpio* B subgenome.

| Category | Content | A/B subgenome |
|---|---|---|
| CID:CCA:CCB=1:0:1 | 1220 | B |
| CID:CCA:CCB=1:1:0 | 915 | A |
| CID:CCA:CCB=1:0:2 total | 110 | B |
| CID:CCA:CCB=1:0:2 oneChr | 88 | B |
| CID:CCA:CCB=1:0:2 twoChr | 22 | B |
| CID:CCA:CCB=1:2:0 total | 126 | A |
| CID:CCA:CCB=1:2:0 oneChr | 102 | A |
| CID:CCA:CCB=1:2:0 twoChr | 24 | A |
| CID:CCA:CCB=1:1:1 total | 8353 | A & B |
| CID:CCA:CCB=1:1:1 paired | 8291 | A & B |
| CID:CCA:CCB=1:1:1 non-paired | 62 | A & B |

**Supplementary Table 16-23. Annotation, GO and KEGG analysis of single copy genes in two subgenomes.**

(Included in a separated excel file)

**Supplementary Table 24-25. Chromsome translocated genes in subgenomes.**

(Included in a separated excel file)

**Supplementary Table 26. Selective pressure of two subgenomes.**

(Included in a separated excel file)

**Supplementary Table 27. Transcriptome data from 12 tissues of *C. carpio*.**

| Tissue | Raw reads | Clean reads | Clean bases (Gbp) | Total mapped | Q20(%) | Q30(%) | GC content(%) |
|---|---|---|---|---|---|---|---|
| Intestine | 58,594,984 | 57,016,106 | 8.55 | 48197574 (84.53%) | 96.6 | 91.48 | 45.81 |
| Liver | 50,842,422 | 40,228,218 | 6.03 | 35166850 (87.42%) | 96.99 | 92.2 | 45.92 |
| Muscle | 46,531,258 | 36,744,052 | 5.51 | 32616748 (88.77%) | 96.8 | 91.72 | 47.79 |
| Brain | 53,510,636 | 42,260,310 | 6.34 | 36590143 (86.58%) | 96.53 | 91.26 | 44.12 |
| Spleen | 49,081,138 | 38,553,610 | 5.78 | 32503886 (84.31%) | 96.01 | 90.22 | 45.27 |
| Skin | 55,044,348 | 43,528,204 | 6.53 | 36769595 (84.47%) | 96.66 | 91.62 | 45.17 |
| Gill | 47,026,858 | 37,045,818 | 5.56 | 31717259 (85.62%) | 96.77 | 91.75 | 44.74 |
| Kidney | 48,014,150 | 37,875,804 | 5.68 | 31945848 (84.34%) | 96.61 | 91.36 | 46.65 |
| Head-kidney | 44,363,242 | 35,042,086 | 5.26 | 29912694 (85.36%) | 96.57 | 91.28 | 47.51 |
| Blood | 38,771,190 | 37,779,062 | 5.67 | 31642491 (83.76%) | 95.08 | 87.7 | 45.4 |
| Sex | 44,232,212 | 34,908,582 | 5.24 | 29941006 (85.77%) | 96.43 | 91.13 | 46.92 |
| Heart | 51,011,754 | 40,348,926 | 6.05 | 35910413 (89%) | 96.62 | 91.35 | 45.72 |

**Supplementary Table 28-51. Annotation of the divergent genes in 12 tissues in *C.carpio*.**

(Included in a separated excel file)

**Supplementary Table 52. Homoeologous gene expression divergence in two subgenomes.**

| Tissues | log2(A/B（FPKM）) | | | | | | | | | |
|---|---|---|---|---|---|---|---|---|---|---|
| | 1 | -1 | 2 | -2 | 3 | -3 | 4 | -4 | 5 | -5 |
| Intestine | 1255 | 1764 | 534 | 837 | 273 | 444 | 147 | 248 | 82 | 148 |
| Liver | 1342 | 1647 | 574 | 774 | 271 | 375 | 132 | 199 | 65 | 104 |
| Muscle | 1439 | 1977 | 577 | 835 | 240 | 341 | 96 | 168 | 46 | 77 |
| Brain | 1258 | 1651 | 447 | 666 | 207 | 308 | 99 | 152 | 53 | 76 |
| Spleen | 1277 | 1696 | 560 | 769 | 274 | 404 | 146 | 202 | 82 | 106 |
| Skin | 1336 | 1800 | 469 | 650 | 178 | 247 | 74 | 87 | 27 | 34 |
| Gill | 1285 | 1822 | 504 | 809 | 231 | 416 | 119 | 216 | 56 | 110 |
| Kidney | 1539 | 1853 | 682 | 852 | 322 | 432 | 165 | 236 | 91 | 130 |
| Head kidney | 1613 | 1945 | 656 | 886 | 281 | 425 | 136 | 223 | 63 | 103 |
| Blood | 1372 | 1643 | 691 | 823 | 375 | 449 | 210 | 257 | 107 | 145 |
| Gonad | 1441 | 1862 | 557 | 822 | 239 | 426 | 114 | 221 | 53 | 113 |
| Heart | 1350 | 1773 | 547 | 819 | 268 | 408 | 130 | 225 | 69 | 112 |
| 12 tissues | 4719 | 5403 | 2590 | 3225 | 1349 | 1802 | 728 | 1067 | 406 | 627 |
| | 62.62% | 71.70% | 51.33% | 63.91% | 45.56% | 60.86% | 41.82% | 61.29% | 39.88% | 61.59% |
| One-way | 2133 | 2817 | 1821 | 2456 | 1159 | 1612 | 674 | 1013 | 391 | 612 |
| | 28.30% | 37.38% | 36.09% | 48.67% | 39.14% | 54.44% | 38.71% | 58.18% | 38.41% | 60.12% |
| Swing | 2586 | | 769 | | 190 | | 54 | | 15 | |
| | 34.32% | | 15.24% | | 6.42% | | 3.10% | | 1.47% | |
| Overall | 7536 | | 5046 | | 2961 | | 1741 | | 1018 | |
| | 91.00% | | 60.93% | | 35.76% | | 21.02% | | 12.29% | |

Note: 8077 of the 8291 homoeologous gene pairs expressed in the 12 tissues.

**Supplementary Table 53-58. Annotation, GO and KEGG analysis of the 32fold-change genes between two subgenomes of the 12 tissues in *C.carpio*.**

(Included in a separated excel file)

**Supplementary Table 59. Transcriptome data from 12 tissues of *C. idella*.**

| Tissue | Raw reads | Clean reads | Clean bases (Gbp) | Total mapped | Q20(%) | Q30(%) | GC content (%) |
|---|---|---|---|---|---|---|---|
| Intestine | 43,258,982 | 41,811,030 | 6.27 | 37306691 (89.23%) | 96.92 | 92.22 | 43.46 |
| Liver | 42,896,884 | 41,453,584 | 6.22 | 37858359 (91.33%) | 96.98 | 92.24 | 45.29 |
| Muscle | 44,991,472 | 43,592,278 | 6.54 | 40341350 (92.54%) | 96.98 | 92.23 | 46.75 |
| Brain | 54,535,354 | 43,120,570 | 6.47 | 39382263 (91.33%) | 96.73 | 91.72 | 43.69 |
| Spleen | 55,518,368 | 43,834,564 | 6.58 | 39105328 (89.21%) | 96.59 | 91.44 | 43.82 |
| Skin | 48,191,282 | 38,073,406 | 5.71 | 34318332 (90.14%) | 96.75 | 91.71 | 45.59 |
| Gill | 48,740,776 | 38,515,054 | 5.78 | 34820462 (90.41%) | 96.66 | 91.56 | 44.01 |
| Kidney | 52,425,680 | 41,487,996 | 6.22 | 37389070 (90.12%) | 96.8 | 91.86 | 43.92 |
| Head-kidney | 47,771,592 | 37,711,358 | 5.66 | 33816587 (89.67%) | 96.63 | 91.48 | 44.23 |
| Blood | 58,843,694 | 57,318,250 | 8.6 | 51253377 (89.42%) | 96.93 | 92.15 | 45.32 |
| Sex | 46,647,678 | 36,735,846 | 5.51 | 32338397 (88.03%) | 96.41 | 91.22 | 46.24 |
| Heart | 50,878,226 | 40,240,458 | 6.04 | 37180686 (92.4%) | 96.79 | 91.74 | 44.77 |

**Supplementary Table 60-61. Annotation of the extremely divergent expressed homoeologous genes.**

(Included in a separated excel file)

**Supplementary Table 62-65. GO and KEGG analysis of the extremely divergent expressed homoeologous genes.**

(Included in a separated excel file)

**Supplementary Table 66. Homoeologous gene expression under abiotic and disease stresses.**

| Stress | Category | Description | NO. of gene pairs |
|---|---|---|---|
| Hypoxia | Expressed gene pairs | FPKM>0 | 6708 |
| | Treatment | A dominant | 1336 |
| | | B dominant | 1870 |
| | | balanced | 3502 |
| | Control | A dominant | 1357 |
| | | B dominant | 1811 |
| | | balanced | 3540 |
| | (A/B)treatment / (A/B)control | A changed faster | 459 |
| | | B changed faster | 528 |
| | | balanced | 5721 |
| CyHV-3 | Expressed gene pairs | FPKM>0 | 7054 |
| | Treatment | A dominant | 1499 |
| | | B dominant | 1982 |
| | | balanced | 3573 |
| | Control | A dominant | 1189 |
| | | B dominant | 1815 |
| | | balanced | 4050 |
| | (A/B)treatment / (A/B)control | A changed faster | 1196 |
| | | B changed faster | 1112 |
| | | balanced | 4746 |
| Aeromonas hydrophila | Expressed gene pairs | FPKM>0 | 6802 |
| | Treatment | A dominant | 1533 |
| | | B dominant | 1938 |
| | | balanced | 3331 |
| | Control | A dominant | 1481 |
| | | B dominant | 1844 |
| | | balanced | 3478 |
| | (A/B)treatment / (A/B)control | A changed faster | 774 |
| | | B changed faster | 831 |
| | | balanced | 5197 |

**Supplementary Table 67. Differentially expressed genes of 8291 homeologous gene pairs under abiotic and biotic stress.**

| Stress | Category | NO. of genes | Ratio_1 | Total_1 | Ratio_2 | Total |
|---|---|---|---|---|---|---|
| Hypoxia | No response | 10980 | 76.53% | | | |
| | Up regulated | 479 | 3.34% | 3367 | 23.47% | 14347 |
| | Down regulated | 2888 | 20.13% | | | |
| CyHV-3 | No response | 9137 | 61.30% | | | |
| | Up regulated | 1584 | 10.63% | 5768 | 38.70% | 14905 |
| | Down regulated | 4184 | 28.07% | | | |
| Aeromonas hydrophila | No response | 11120 | 76.33% | | | |
| | Up regulated | 1533 | 10.52% | 3448 | 23.67% | 14568 |
| | Down regulated | 1915 | 13.15% | | | |

**Supplementary Table 68. Differentially expressed homeologous gene pairs of the 8291 homeologous gene pairs under abiotic and biotic stress.**

| Stress | Category | NO. of gene pairs | Ratio_1 | Total_1 | Ratio_2 | Total |
|---|---|---|---|---|---|---|
| Hypoxia | No response | 5553 | 82.78% | | | |
| | Up regulated | 126 | 1.88% | 1155 | 17.22% | 6708 |
| | Down regulated | 1029 | 15.34% | | | |
| CyHV-3 | No response | 4757 | 67.44% | | | |
| | Up regulated | 613 | 8.69% | 2297 | 32.56% | 7054 |
| | Down regulated | 1684 | 23.87% | | | |
| Aeromonas hydrophila | No response | 5615 | 82.55% | | | |
| | Up regulated | 607 | 8.92% | 1187 | 17.45% | 6802 |
| | Down regulated | 580 | 8.53% | | | |

**Note:** Homoeologous pairs had Treatment (A+B) expression values 2-fold less than Control (A+B) expression during three stress treatments were regarded as no response.

**Supplementary Table 69. Whole genome methylation levels of common carp.**

| Samples | mC (%) | mCpG (%) | mCHG | mCHH |
|---|---|---|---|---|
| 01 | 5.74% | 64.99% | 0.17% | 0.39% |
| 02 | 5.83% | 65.64% | 0.17% | 0.45% |
| 03 | 5.89% | 66.32% | 0.18% | 0.45% |

**Supplementary Table 70. Whole genome resequencing of YR and HB.**

| Sample name | Raw base | Q20(%) | Q30(%) | GC content(%) | Clean reads | Mapped reads | Mapping rate | Average depth |
|---|---|---|---|---|---|---|---|---|
| HB1 | 29,728,057,800 | 97.49 | 94.58 | 37.73 | 172,613,220 | 169,891,480 | 98.42% | 17.64 |
| HB2 | 31,813,682,700 | 97.43 | 94.41 | 37.83 | 186,948,806 | 183,242,802 | 98.02% | 18.82 |
| HB3 | 30,394,545,000 | 97.51 | 94.53 | 38.13 | 182,321,940 | 179,318,793 | 98.35% | 18.54 |
| HB4 | 26,683,970,400 | 97.54 | 94.55 | 37.98 | 159,576,502 | 156,975,115 | 98.37% | 16.22 |
| HB5 | 26,623,073,100 | 97.80 | 95.09 | 37.95 | 161,142,628 | 158,701,832 | 98.49% | 16.63 |
| HB6 | 30,735,050,700 | 97.66 | 94.84 | 38.05 | 186,318,212 | 183,210,710 | 98.33% | 19.1 |
| HB7 | 30,526,950,000 | 97.13 | 93.70 | 37.99 | 173,959,922 | 171,137,221 | 98.38% | 17.54 |
| HB8 | 28,422,233,100 | 97.02 | 93.48 | 38.11 | 162,113,712 | 159,274,297 | 98.25% | 16.28 |
| HB9 | 28,514,219,100 | 97.26 | 94.02 | 37.98 | 159,128,022 | 156,354,487 | 98.26% | 15.91 |
| HB10 | 30,851,652,000 | 97.52 | 94.44 | 38.04 | 167,269,534 | 164,428,446 | 98.30% | 16.52 |
| HB11 | 30,663,478,200 | 97.73 | 94.99 | 38.31 | 157,789,474 | 154,844,387 | 98.13% | 15.79 |
| HB12 | 28,604,481,300 | 97.70 | 94.89 | 38.19 | 148,817,324 | 146,149,883 | 98.21% | 14.98 |
| HB13 | 25,777,352,700 | 97.54 | 94.49 | 38.10 | 132,002,382 | 129,563,204 | 98.15% | 13.04 |
| HB14 | 28,788,098,700 | 97.52 | 94.48 | 37.99 | 145,352,438 | 142,698,022 | 98.17% | 14.28 |
| HB15 | 31,686,123,300 | 97.54 | 94.51 | 37.96 | 160,391,024 | 157,305,510 | 98.08% | 15.71 |
| HB17 | 28,814,374,500 | 97.66 | 94.81 | 38.29 | 157,173,936 | 154,514,115 | 98.31% | 15.8 |
| HB18 | 31,957,320,600 | 97.56 | 94.58 | 38.19 | 175,749,860 | 172,818,825 | 98.33% | 17.68 |
| HB19 | 29,366,688,300 | 97.73 | 94.91 | 37.92 | 162,011,046 | 159,595,585 | 98.51% | 16.49 |
| YR01 | 26,740,550,700 | 97.50 | 94.66 | 38.05 | 160,432,162 | 157,404,584 | 98.11% | 16.38 |
| YR02 | 29,289,688,500 | 97.49 | 94.59 | 37.90 | 171,367,532 | 168,498,411 | 98.33% | 17.38 |
| YR03 | 24,784,188,600 | 97.53 | 94.73 | 38.04 | 148,910,426 | 146,345,250 | 98.28% | 15.21 |
| YR04 | 22,500,513,000 | 97.31 | 94.28 | 38.03 | 144,009,038 | 141,261,876 | 98.09% | 14.47 |
| YR05 | 29,557,388,700 | 97.53 | 94.61 | 38.17 | 154,482,400 | 151,436,217 | 98.03% | 15.3 |
| YR06 | 27,444,222,600 | 97.25 | 93.92 | 37.91 | 152,029,730 | 149,332,295 | 98.23% | 14.9 |
| YR07 | 29,574,770,400 | 97.37 | 94.33 | 38.00 | 179,459,908 | 176,416,175 | 98.30% | 18.31 |
| YR08 | 28,798,885,800 | 97.49 | 94.50 | 37.95 | 175,002,800 | 172,154,966 | 98.37% | 18.04 |

| | | | | | | | | |
|---|---|---|---|---|---|---|---|---|
| YR09 | 28,474,039,800 | 97.59 | 94.71 | 38.06 | 156,216,146 | 153,538,743 | 98.29% | 15.52 |
| YR10 | 29,111,964,000 | 97.34 | 94.07 | 38.04 | 163,129,036 | 160,592,272 | 98.44% | 16.27 |
| YR11 | 26,988,921,900 | 97.14 | 93.74 | 37.93 | 152,677,662 | 150,213,207 | 98.39% | 15.23 |
| YR12 | 34,672,018,200 | 98.28 | 96.23 | 37.92 | 182,982,352 | 179,217,538 | 97.94% | 17.89 |
| YR13 | 31,183,845,600 | 98.35 | 96.37 | 37.65 | 164,943,872 | 161,603,412 | 97.97% | 16.17 |
| YR14 | 28,855,017,000 | 96.59 | 92.58 | 37.85 | 176,341,922 | 172,835,544 | 98.01% | 17.77 |
| YR15 | 25,900,995,000 | 96.38 | 92.16 | 37.70 | 158,263,722 | 155,193,174 | 98.06% | 16.09 |
| YR16 | 29,124,218,700 | 95.44 | 90.59 | 38.04 | 179,997,668 | 175,547,085 | 97.53% | 17.9 |

**Supplementary Table 71. Selection signatures of HB and YR.**

| Strains | genomic regions | Selective size (bp) | Selective genes |
|---|---|---|---|
| HB | 248 | 29,725,000 | 1237 |
| YR | 120 | 11,075,000 | 470 |
